# Cohesin–axis interaction via a conserved Red1 motif promotes domain-specific DSB formation and Mek1 activation

**DOI:** 10.64898/2026.07.03.736430

**Authors:** Arrosan Rajalingam, Yusuke Tsuruta, Taniya Roy, Julian Urdiain-Arraiza, Hasan F Alnaser, Shin-ichiro Hiraga, Corentin Claeys Bouuaert, Hajime Murakami

## Abstract

Faithful chromosome segregation during meiosis I requires tight control of interhomolog recombination. In budding yeast, the meiotic chromosome axis, built on Rec8-containing cohesin together with Red1 and Hop1, acts as a central platform regulating meiotic recombination from programmed DNA double-strand break (DSB) formation to checkpoint signaling and chromosome segregation, yet how cohesin recruits axis proteins remains unclear. Here, we identified a conserved cohesin-interacting motif (CIM) in Red1 that directly binds Rec8. AlphaFold3 modeling predicted that Red1-CIM forms a short α-helix that docks into a conserved hydrophobic pocket within the Rec8 C-terminal winged-helix domain, which we confirmed biochemically. Disruption of the Red1-CIM preferentially impaired Red1 recruitment to Rec8-dependent chromosomal regions, while relative enrichment in Rec8-independent domains was preserved, leading to reduced DSB formation in Rec8-dependent domains. The Red1-CIM mutation also reduced crossover formation, increased chromosome missegregation, and reduced spore viability. Notably, this spore lethality exceeded that predicted by the reduction in DSB formation. Consistently, *red1-CIM* mutants failed to activate the meiotic checkpoint kinase Mek1. Finally, we provide evolutionary, structural, and biochemical evidence that this Red1–Rec8 interaction is conserved across fungi and plants. Together, these findings define a direct molecular bridge linking cohesin to chromosome-axis organization, spatial DSB regulation, and checkpoint signaling during meiosis.

## Introduction

Meiotic recombination generates genetic diversity and ensures faithful chromosome segregation during gametogenesis (Borner et al. 2023; Zickler and Kleckner 2023). During meiotic prophase, programmed DNA double-strand breaks (DSBs) introduced by the conserved Spo11 transesterase and its essential accessory factors (DSB proteins) initiate homologous recombination, ultimately yielding interhomolog crossovers that physically link homologous chromosomes and promote their accurate segregation at the first meiotic division (Lam and Keeney 2014; Yadav and Claeys Bouuaert 2021). Because errors in DSB formation or repair compromise chromosome segregation and gamete viability, both the initiation and progression of meiotic recombination are subject to stringent spatial and temporal regulation.

A major platform for implementing this regulation is the meiotic chromosome axis, a meiosis-specific proteinaceous structure that organizes chromatin into arrays of loops anchored at their bases along a linear axial core (Zickler and Kleckner 1999; Ur and Corbett 2021). In budding yeast, this axis is built upon cohesin complexes containing the meiosis-specific kleisin Rec8, together with the axis proteins Red1 and Hop1 (Hollingsworth et al. 1990; Smith and Roeder 1997; Klein et al. 1999; Panizza et al. 2011). Rec8-containing cohesin establishes sister chromatid cohesion and may also promote loop extrusion, thereby providing the structural framework for loop–axis architecture (Klein et al. 1999; Schalbetter et al. 2019). Red1 and Hop1 assemble on this cohesin-based scaffold throughout meiotic prophase and constitute regulatory components of the meiotic chromosome axis (Humphryes and Hochwagen 2014; Subramanian and Hochwagen 2014).

Chromosome axis proteins promote meiotic DSB formation. At the scale of chromosomal domains (>20 kb), DSB enrichment correlates with local Hop1 and Red1 levels (Blat et al. 2002; Pan et al. 2011; Panizza et al. 2011; Sun et al. 2015). Loss of axis proteins reduces DSB formation at multiple DSB hotspots and chromosomal regions (Blat et al. 2002; Kugou et al. 2009; Kim et al. 2010), although the extent of reduction varies depending on the locus examined, with *hop1* showing the strongest overall defect (Mao-Draayer et al. 1996; Xu et al. 1997; Woltering et al. 2000; Pecina et al. 2002; Niu et al. 2005). Hop1 promotes DSB formation through a conserved interaction with Mer2, facilitating assembly of the DSB machinery on chromosome axes (Acquaviva et al. 2013; Sommermeyer et al. 2013; Stanzione et al. 2016; Kariyazono et al. 2019; Rousova et al. 2021). Consistently, artificial targeting of Hop1 to normally DSB-poor regions creates ectopic DSB hotspots (Shodhan et al. 2022), supporting its role as a primary axis component promoting DSB competence.

Following DSB formation, Red1 and Hop1 play central roles in activating the meiosis-specific kinase Mek1, a Rad53/CHK2 paralog. Mek1 is recruited to chromosome axes and activated via Mec1/Tel1-dependent phosphorylation of Hop1 in response to Spo11-induced DSBs (Hollingsworth et al. 1990; Smith and Roeder 1997; Niu et al. 2005; Carballo et al. 2008).

Activated Mek1 enforces interhomolog recombination bias by suppressing sister-chromatid repair, thereby promoting crossover formation between homologs (Schwacha and Kleckner 1997; Niu et al. 2009; Kim et al. 2010). Accordingly, *red1, hop1*, and *mek1* mutants exhibit severe chromosome segregation defects and drastically reduced spore viability due to meiosis I nondisjunction (Rockmill and Roeder 1988; Hollingsworth and Byers 1989; Rockmill and Roeder 1991).

Red1 contains an N-terminal domain composed of Armadillo (ARM) and Pleckstrin Homology (PH) subdomains, followed by a long intrinsically disordered region (IDR) and a C-terminal coiled-coil (CC) domain (**Fig. 1A**). The CC domain forms parallel–antiparallel homotetramers that assemble end-to-end into filaments *in vitro*, which is proposed to underlie the core repeating unit of the meiotic axis (West et al. 2019). Adjacent to the N-terminal domain, Red1 contains a closure motif (CM) that binds the HORMA domain of Hop1, thereby enabling Hop1 recruitment to the axis (Woltering et al. 2000; West et al. 2018). Hop1 also contains a central chromatin-binding region (CBR) implicated in nucleosome association (Milano et al. 2024) and a C-terminal closure motif that mediates Hop1 homo-oligomerization (Niu et al. 2005; West et al. 2018). Cohesin architecture is likewise well defined: the mitotic kleisin Scc1 binds the Smc3 coiled-coil neck region via its N terminus and the Smc1 head via its C-terminal winged-helix domain, while its central region associates with HEAT-repeat proteins (Pds5, Scc2, and Scc3) (Gligoris and Lowe 2016).

**Figure 1.**
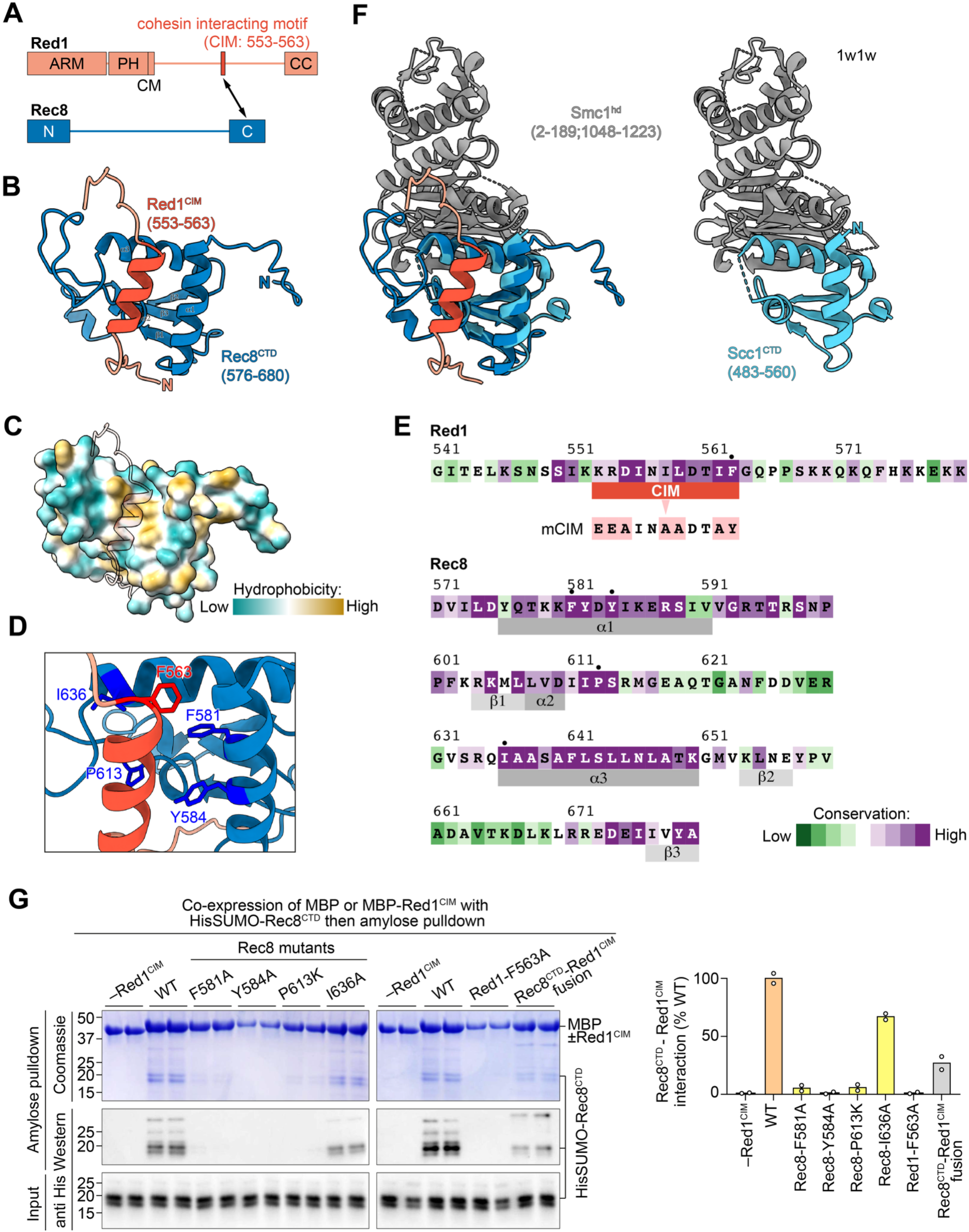
A conserved Red1 motif interacts with the Rec8 C-terminal domain. **A.** Domain organization of Red1 and Rec8. Red1 contains an armadillo-like domain (ARM; residues 1–225), a pleckstrin homology–like domain (PH; residues 232–344), a closure motif (CM; residues 345–362), and a C-terminal coiled-coil domain (CC; residues 734–827). A putative cohesin-interacting motif (CIM; residues 553–563) is located between the CM and CC domains and is predicted to interact with the Rec8 C-terminal domain (CTD; residues 576–680). **B.** AlphaFold3 prediction of the Red1-CIM (red) bound to the Rec8-CTD (blue). **C.** Surface representation of the predicted Rec8 CTD highlighting hydrophobicity. The Red1-CIM (semi-transparent cartoon) is shown bound within a hydrophobic pocket. **D.** Close-up view of the predicted interface showing key residues on both Red1 and Rec8. **E.** Sequence conservation of the Red1-CIM and the Rec8 CTD interface across Saccharomycetaceae species. Amino acid substitutions introduced in yeast are indicated below the CIM. Dots indicate residues tested in panel G. **F.** Structural comparison showing that the Red1-CIM binding site on the Rec8-CTD is spatially distinct from the interface between the kleisin and the Smc1 head domain. The crystal structure of the Scc1-CTD bound to the Smc1 head domain (right; PDB 1W1W) is overlaid with the AlphaFold model of the Rec8-CTD bound to the Red1-CIM (left). **G.** *In vitro* binding assays. MBP or MBP–Red1 fragments were co-expressed with HisSUMO–Rec8-CTD, followed by amylose pulldown. Mutations in Rec8 or Red1 reduce the interaction. Right: quantification of pulldown efficiency relative to wild type. Bars represent the mean from two independent experiments (open circles).

Rec8 likely preserves a similar overall domain organization (Gruber et al. 2003; Schleiffer et al. 2003), suggesting conservation of core cohesin assembly principles (Sakuno and Hiraoka 2022).

Loss of Rec8 globally reduces chromosomal Red1 and Hop1 association, except at distinct genomic regions that recruit prominent Red1–Hop1, defining Rec8-dependent and independent pathways recruiting axis proteins (Panizza et al. 2011; Sun et al. 2015). In “islands,” Hop1 associates with nucleosomes via its chromatin-binding region (CBR), recruiting both Hop1 and Red1 independently of Rec8 (Heldrich et al. 2022; Milano et al. 2024). In contrast, “deserts” require Rec8 for Red1–Hop1 accumulation. Notably, expression of the mitotic kleisin Scc1 in meiotic *rec8Δ* cells restores sister chromatid cohesion but fails to rescue Red1 and Hop1 recruitment in deserts, demonstrating a meiosis-specific role for Rec8 beyond sister chromatid cohesion (Toth et al. 2000; Brar et al. 2009; Sun et al. 2015). Consistently, co-immunoprecipitation and proximity-labeling experiments have revealed association of Rec8 with Red1–Hop1 (Sun et al. 2015). However, whether they interact directly—and through which domains—remains unknown.

Here, we identify a conserved motif in Red1 that directly interacts with the C-terminal domain (CTD) of Rec8. Disruption of this cohesin-interacting motif (CIM) of Red1 selectively impairs Rec8-dependent recruitment of Red1 to meiotic chromosomes and reduces DSB formation preferentially within Rec8-dependent chromosomal domains. Strikingly, despite retaining substantial levels of total DSBs, *red1-CIM* mutants exhibit severe post-DSB defects, indicating that the association of Red1 with cohesin is important for Mek1 activation and faithful meiotic recombination.

## Results

### A conserved Red1 motif interacts with the C-terminal domain of Rec8

To understand the molecular basis of Red1 recruitment by cohesin, we tested the hypothesis that the cohesin subunit Rec8 directly interacts with Red1. Using AlphaFold3, we identified a putative interaction between a short motif of Red1 (residues 553–563) located within its disordered region and the C-terminal domain (CTD) of Rec8 (**Fig. 1A,B**; **Supplemental Fig. S1A**). The respective domains were consistently predicted to interact when the full protein sequences were used as input. When the modeling was performed with the relevant sequences alone, AlphaFold3 yielded moderate-confidence models (global pTM 0.67, ipTM 0.65, **Supplemental Fig. S1B**), supporting a direct association. We therefore refer to this Red1 segment (residues 553–563) as a putative cohesin-interacting motif (CIM).

In the predicted structure, the Red1-CIM folds into an α-helix that engages a hydrophobic pocket within the winged-helix domain of the Rec8-CTD (**Fig. 1C,D**). Interface residues on both Red1 and Rec8 are conserved across representative Saccharomycetaceae species (**Fig. 1E**; **Supplemental Fig. S1C,D**; broader conservation is examined below). This surface is distinct from that of mitotic kleisin Scc1 (**Fig. 1F**; **Supplemental Fig. S1E**), and, consistently, AlphaFold does not predict an interaction between the Red1 peptide and Scc1 (**Supplemental Fig. S1F**). Importantly, the predicted Red1-binding surface on Rec8-CTD is spatially distinct from the interface through which Rec8 engages the Smc1 head domain (**Fig. 1F**) (Haering et al. 2004), suggesting that Red1-Rec8 interaction would be compatible with cohesin ring assembly.

To test this putative interaction between Red1 and Rec8, we co-expressed HisSUMO-tagged Rec8-CTD in *E. coli* with either MBP alone or MBP fused to the Red1 peptide, and tested the interaction by amylose pulldown. MBP-Red1 efficiently pulled down HisSUMO-Rec8-CTD, whereas MBP alone did not, consistent with the interaction predicted by AlphaFold (**Fig. 1G**). In addition, substitutions of Rec8 interface residues F581A, Y584A, and P613K strongly reduced the interaction, while I636A had a more modest effect (**Fig. 1G**).

Reciprocally, mutation of a conserved phenylalanine within Red1 (F563A) markedly reduced binding to Rec8-CTD (**Fig. 1G**). Furthermore, fusing Rec8-CTD to the Red1 motif reduced the interaction with MBP-Red1, indicating that the fused peptide effectively acts as a competitor. Across samples, input levels of Rec8-CTD were comparable, indicating that the reduced pulldown reflected impaired binding rather than differences in protein abundance.

Together, these data demonstrate that Red1 residues 553–563 function as a cohesin-interacting motif (Red1-CIM), suggesting that this interaction may promote Red1 recruitment during meiosis.

### Mutations in Red1-CIM selectively impair Rec8-dependent Red1 recruitment

To test the functional relevance of the Red1–Rec8 interaction, we generated two strains in which the CIM coding sequence at the endogenous *RED1* locus was either mutated (*red1-mCIM*, **Fig. 1E**) or deleted (*red1-ΔCIM*). Both mutants showed similar meiotic progression to wild type (**Supplemental Fig. S2A**). When introduced into a V5-tagged Red1 background, mutant protein levels were comparable to wild-type V5-Red1 (**Supplemental Fig. S2B**).

However, the *red1-ΔCIM* mutant showed reduced nuclear localization of Red1 (**Supplemental Fig. S2C,D**), possibly due to disruption of a putative nuclear localization signal equivalent to that reported in fission yeast Rec10 (Wintrebert et al. 2021) (**Supplemental Fig. S2E**). In contrast, the *red1-mCIM* mutant localized Red1 at a level comparable to wild type (**Supplemental Fig. S2C,D**). Because mutations on the Rec8-CTD reduced Rec8 protein abundance *in vivo* (**Supplemental Fig. S2F**), we focused subsequent functional analyses on Red1 CIM alleles.

We next assessed Red1 chromatin association by V5-Red1 ChIP-seq using cells collected 3 h into meiosis. As a reference condition in which Rec8-dependent Red1 recruitment is impaired, we used a strain that expresses Scc1 during meiosis instead of Rec8 (*pREC8-SCC1*), which preserves cohesion while disrupting Rec8-dependent Red1 recruitment (Toth et al. 2000; Sun et al. 2015). To enable quantitative comparisons across datasets, we used an *S. mikatae* strain expressing Rad50-V5 as a spike-in control (see Methods). Total calibrated Red1 ChIP signal was substantially higher in all V5-tagged strains than the untagged control, while *pREC8-SCC1*, *red1-mCIM*, and *red1-ΔCIM* showed reduced total ChIP signal levels (71, 57, and 51% of wild type, **Fig. 2A**). Biological replicates were highly reproducible (**Supplemental Fig. S3A**), and replicate datasets were therefore averaged for downstream analyses.

**Figure 2.**
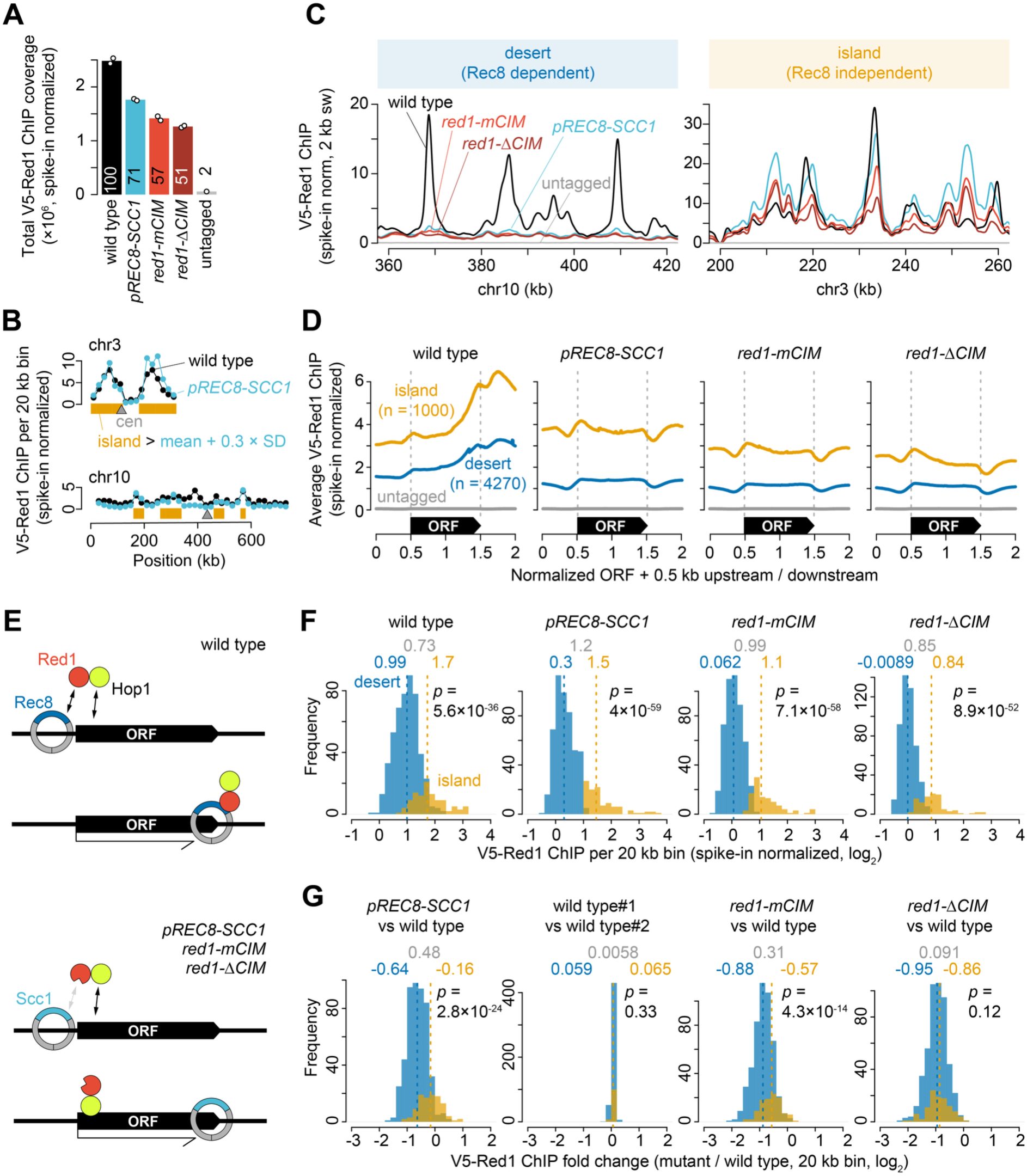
Red1-CIM is required for Rec8-dependent recruitment of Red1 to meiotic chromosomes. **A.** Total calibrated V5-Red1 ChIP-seq signal in wild type, *pREC8-SCC1*, *red1-mCIM*, and the untagged control strain. The untagged control was an untagged *red1-ΔCIM* strain used to estimate nonspecific anti-V5 ChIP background. Bars indicate the mean of biological replicates (open circles). Numbers within bars indicate the percentage of the wild-type mean. **B.** Rec8-independent “island” calling. Red1 ChIP signals in *pREC8-SCC1* were binned into 20-kb windows. Bins showing Red1 ChIP signals above the mean + 0.3 standard deviation (SD) were called islands, and the remaining bins were classified as deserts. With this classification, 17.5% of bins (107/611) were called islands (see also **Supplemental Fig. S3B**). **C.** Representative V5-Red1 ChIP-seq profiles in Rec8-dependent desert and Rec8-independent island regions. Profiles were smoothed with a 2-kb Parzen (triangular) sliding window. **D.** Distribution of Red1 relative to open reading frames (ORFs). Using an R package provided by the Hochwagen laboratory (Sun et al. 2015), ORFs longer than 500 bp were standardized to a length of 1 kb, then ChIP-seq profiles were averaged over the standardized ORFs plus 0.5 kb of upstream and downstream sequence. n indicates the number of ORFs included in the analysis. **E.** Model for Red1/Hop1 relocation as cohesin is displaced by transcription. When the Red1–cohesin interaction is impaired, Red1/Hop1 remain closer to their original recruitment sites. **F.** Genome-wide distribution of V5-Red1 ChIP signal in 20-kb bins in Rec8-dependent deserts and Rec8-independent islands. In F and G, and in Fig. 3D**,E**, dashed lines and numbers indicate medians (grey numbers indicate median differences). *P* values were calculated using two-sided Wilcoxon tests comparing island and desert bins. **G.** Distribution of log2 fold changes (mutant/wild type) in V5-Red1 ChIP signal across 20-kb bins. The wild-type replicate comparison serves as a negative control for fold-change analysis and shows little systematic difference between replicate datasets.

To examine Red1 distribution at chromosomal domain resolution, we binned ChIP signals in 20-kb windows and examined Red1 profiles along all 16 chromosomes (**Supplemental Fig. S3B–D**). Consistent with previous studies (Sun et al. 2015)*, pREC8-SCC1* caused a broad reduction in Red1 binding, although Red1 signal was retained in discrete chromosomal regions, including many regions previously classified as Rec8-independent Red1 recruitment domains, or “islands” (Heldrich et al. 2022). We therefore classified 20-kb bins into islands and Rec8-dependent “deserts” using the *pREC8-SCC1* Red1 ChIP signal as an operational criterion (**Fig. 2B**; see Discussion). In representative genomic regions, *red1-mCIM* reduced Red1 binding most strongly in deserts, whereas substantial Red1 signal was retained in islands. In a desert region on chr10, prominent Red1 peaks observed in wild type were largely diminished in *pREC8-SCC1*, *red1-mCIM*, and *red1-ΔCIM* (**Fig. 2C**, left). In contrast, in an island region on chr3, substantial Red1 signal was retained, particularly in *pREC8-SCC1* and *red1-mCIM*, although the total signal was reduced and relative peak intensities differed from the wild type (**Fig. 2C**, right).

Metagene analysis around ORFs further revealed a change in the spatial pattern of Red1 binding. In wild type, Red1 showed a pronounced enrichment near transcription termination sites (TTSs) (**Fig. 2D**). This TTS-proximal enrichment was largely lost in *pREC8-SCC1* and in the *red1-CIM* mutants, resulting in flatter profiles around ORFs. This pattern is consistent with the idea that cohesin accumulates near TTSs as it is repositioned by transcription, as previously proposed (Sun et al. 2015), and suggests that Red1 enrichment at these sites depends, at least in part, on its interaction with Rec8 (**Fig. 2E**; see Discussion).

We next quantified the impact of disrupting the Red1-CIM on its genome-wide distribution using 20-kb bins. In wild type, Red1 ChIP signal was significantly higher in islands than in deserts, with island bins showing ∼1.7-fold higher median signal than desert bins (**Fig. 2F**). This relative enrichment was retained in *pREC8-SCC1* (∼2.3-fold), in *red1-mCIM* (∼2.0-fold), and *red1-ΔCIM* (∼1.8-fold), consistent with Rec8-independent recruitment of Red1 in islands (**Fig. 2F**).

Relative to wild type, *pREC8-SCC1* and *red1-mCIM* showed stronger reductions in Red1 signal in deserts than in islands, consistent with a role for Rec8 in Red1 recruitment in deserts (**Fig. 2G**). In *pREC8-SCC1*, median Red1 signal was reduced to ∼64% of wild type in deserts but retained at ∼90% in islands; similarly, in *red1-mCIM*, it was reduced to ∼54% in deserts and ∼67% in islands. By contrast, *red1-ΔCIM* showed similar reductions in deserts and islands (∼52% and ∼55% of wild type, respectively), which may be a consequence of the reduced nuclear localization of Red1 in this strain (**Supplemental Fig. S2C,D**; see Discussion).

Together, these analyses indicate that the interaction between Red1 and Rec8 drives Red1 recruitment within deserts and controls Red1 localization within islands, where it is recruited largely independently of Rec8.

### Red1-CIM mutation reduces DSB formation in Rec8-dependent chromosomal domains

To assess how impaired Red1 recruitment affects meiotic DSB formation, we performed S1-seq in *sae2Δ*, *red1-mCIM sae2Δ,* and *red1Δ sae2Δ* strains harvested 3.5 h after meiotic induction, and in *sae2Δ* and *red1-mCIM sae2Δ* cells harvested at 6 h. To enable quantitative comparisons across samples, datasets were calibrated using *S. mikatae sae2*Δ spike-in cells collected 6 h into meiosis (see Methods; **Fig. 3A**).

**Figure 3.**
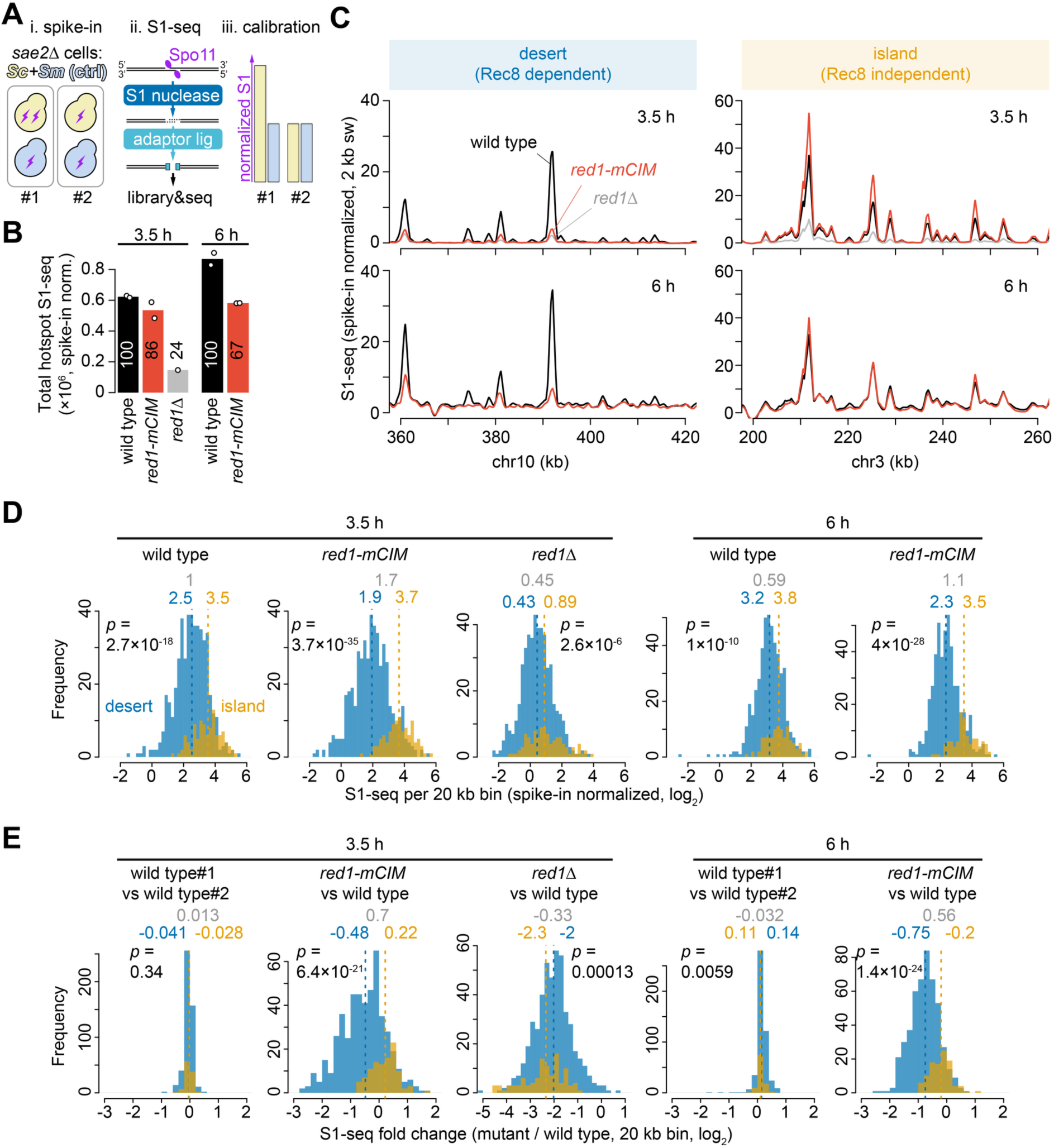
S1-seq reveals region-specific effects of Red1-CIM mutation on meiotic DSB formation. **A.** Schematic of the spike-in calibrated S1-seq workflow. Meiotic *S. cerevisiae* (*Sc*) samples were mixed with a fixed number of *S. mikatae* (*Sm*) cells as a spike-in control, treated with S1 nuclease to process Spo11-associated DSB ends, and subjected to adaptor ligation, library preparation, and deep sequencing. Signals were calibrated between samples using the spike-in control. **B.** Total hotspot-associated S1-seq signal within ±100 bp of Spo11-oligo hotspot peaks (Mohibullah and Keeney 2017). The same hotspot set was used for all genotypes and time points. Signals are shown for wild type, *red1-mCIM*, and *red1Δ* at 3.5 h, and for wild type and *red1-mCIM* at 6 h after meiotic induction, after spike-in normalization. Bars indicate the mean of biological replicates (open circles). Numbers within bars indicate the percentage of the wild-type mean. **C.** Representative S1-seq profiles in Rec8-dependent desert and Rec8-independent island regions. Profiles were smoothed with a 2-kb Parzen (triangular) sliding window. **D.** Genome-wide distribution of S1-seq signal in 20-kb bins in Rec8-dependent deserts and Rec8-independent islands. **E.** Distribution of S1-seq fold changes relative to wild type in 20-kb bins. Log2 mutant/wild-type ratios are shown for deserts and islands. The wild-type replicate comparisons serve as negative controls for fold-change analysis and show little systematic difference between replicate datasets.

We quantified S1-seq signal using a published Spo11-oligo hotspot annotation (Mohibullah and Keeney 2017), providing a common set of DSB sites for comparison across genotypes and time points. Total hotspot-associated S1-seq signal was reduced in *red1-mCIM* relative to wild type at both time points, to 86% at 3.5 h and 67% at 6 h of the corresponding wild-type level, whereas *red1Δ* showed a much stronger reduction (to 24% of wild type) at 3.5 h (**Fig. 3B**). Biological replicates showed high reproducibility (**Supplemental Fig. S4A**), and replicate datasets were therefore averaged for downstream analyses. Our datasets were in good agreement with previously published *sae2Δ* S1-seq data (Huang et al. 2025) and Spo11-oligo map (Mohibullah and Keeney 2017) (**Supplemental Fig. S4B,C**).

We next examined whether the reduction in DSB signal was biased toward specific chromosomal domains. Using the same island/desert classification defined from Red1 ChIP data, we compared S1-seq patterns in representative genomic regions. In a Rec8-dependent desert region on chr10, S1-seq signal was clearly reduced in *red1-mCIM* relative to wild type at both 3.5 h and 6 h, whereas in an island region on chr3 the effect was much weaker and DSB signal was largely retained (**Fig. 3C**). As expected, *red1Δ* caused a strong reduction in S1-seq signal, including in representative desert and island regions (**Fig. 3C**). This overall tendency was also apparent in chromosome-scale profiles across all 16 chromosomes (**Supplemental Fig. S4D–F**).

To quantify this domain bias genome-wide, we compared the distributions of S1-seq signal across 20-kb bins. In wild type, S1-seq signal was higher in islands than in deserts at both 3.5 h and 6 h, with island/desert median ratios of ∼2.0-fold and ∼1.5-fold, respectively (**Fig. 3D**). In *red1-mCIM*, this separation increased to ∼3.2-fold at 3.5 h and ∼2.1-fold at 6 h, consistent with preferential reduction of breaks in deserts. In contrast, in *red1Δ*, the two distributions became more similar, with an island/desert ratio of ∼1.4-fold at 3.5 h, consistent with reduced S1-seq signal in both islands and deserts (**Fig. 3D**). These conclusions were confirmed by comparing changes in S1-seq signal relative to wild type. At 3.5 h, *red1-mCIM* reduced median S1-seq signal in deserts to ∼72% of wild type, whereas island signal was largely retained at ∼117% (**Fig. 3E**). At 6 h, a similar but weaker domain bias was evident, with desert and island signals reduced to ∼59% and ∼87% of wild type, respectively. In contrast, *red1Δ* caused a strong reduction in both deserts and islands, to ∼25% and ∼20% of wild type, respectively (**Fig. 3E**).

Together, these results indicate that disruption of the Red1–Rec8 interaction significantly compromises DSB formation in regions where Red1 recruitment depends on Rec8, leading to modestly reduced global DSB levels.

### Loss of Red1 binding in *red1-mCIM* predicts reduced DSB formation in Rec8-dependent domains

We next directly compared the effect of the *red1-mCIM* mutation on Red1 chromatin association and DSB formation. In representative genomic regions, reductions in Red1 ChIP signal broadly paralleled reductions in S1-seq signal. For example, in desert regions where Red1 binding was strongly reduced in *red1-mCIM*, S1-seq signal was also reduced, whereas island regions with relatively preserved Red1 binding showed much smaller changes in DSB signal (**Fig. 4A**). This tendency was also apparent in chromosome-scale fold-change profiles, where reductions in S1-seq signal were most evident in regions showing reduced Red1 ChIP, particularly in Rec8-dependent desert regions (**Fig. 4B**; **Supplemental Fig. S5A**).

**Figure 4.**
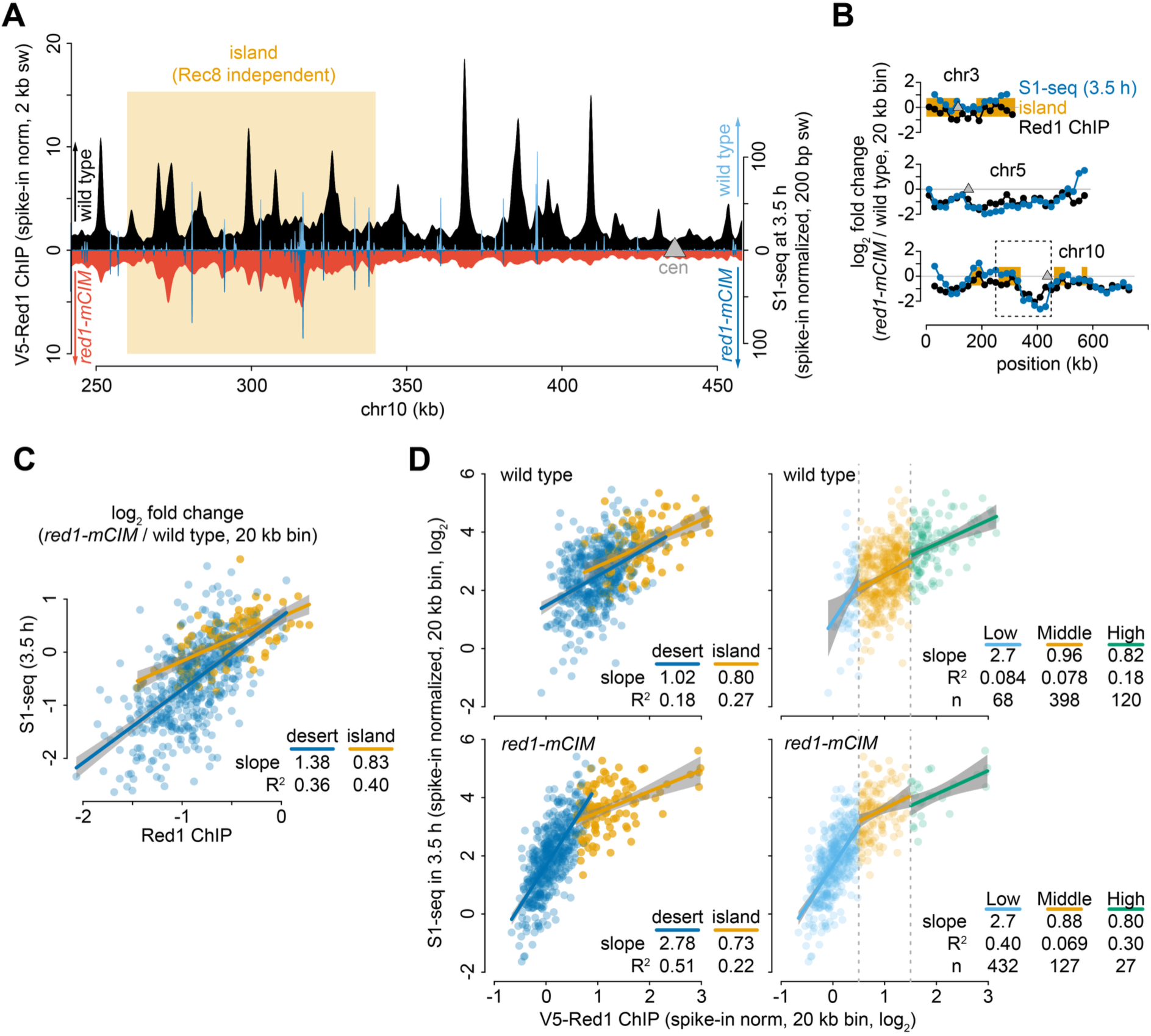
Red1-CIM-dependent changes in Red1 chromosomal association are coupled to changes in DSB formation. **A.** Representative profiles of spike-in calibrated V5-Red1 ChIP-seq and S1-seq signals across a chr10 interval in wild type and *red1-mCIM*. V5-Red1 ChIP-seq signals are shown in black and red and were smoothed with a 2-kb Parzen sliding window. S1-seq signals are shown in light blue and blue and were smoothed with a 200-bp Parzen sliding window. For *red1-mCIM*, the y-axis is inverted, so stronger signals extend downward. **B.** Representative profiles of log2 fold changes in V5-Red1 ChIP-seq and S1-seq signals on chr3, chr5, and chr10 in *red1-mCIM* relative to wild type. Dashed box in chr10 indicates the region shown in panel A. **C.** Relationship between changes in V5-Red1 ChIP-seq and S1-seq signals in *red1-mCIM* relative to wild type. Each point represents a 20-kb bin. Rec8-dependent deserts and Rec8-independent islands are shown separately. In panels C and D, lines indicate linear regression fits; shaded bands indicate 95% confidence intervals for the fitted mean. Slopes and R^2^ values are indicated. **D.** Relationship between V5-Red1 ChIP-seq signal and S1-seq signal at 3.5 h in wild type and *red1-mCIM*. Calibrated signals in 20-kb bins are log2-transformed. Left panels show bins classified as Rec8-dependent deserts or Rec8-independent islands. Right panels show the same data stratified by V5-Red1 ChIP signal into low, middle, and high Red1 bins (break points: 0.5 and 1.5). Slopes, R^2^ values, and numbers of bins are indicated.

To quantify this relationship genome-wide, we compared fold changes in Red1 ChIP and S1-seq signal in *red1-mCIM* relative to wild type across 20-kb bins. Reductions in Red1 ChIP were positively associated with reductions in S1-seq signal, indicating that the local impact of *red1-mCIM* on DSB formation broadly follows its impact on Red1 chromatin association (**Fig. 4C**). This relationship was observed in both deserts and islands, but was steeper in deserts, consistent with DSB formation in Rec8-dependent domains being more sensitive to reduced Red1 binding.

We further examined the relationship between absolute Red1 ChIP signal and S1-seq signal. In wild type, Red1 ChIP and S1-seq were positively associated in both deserts and islands, although the relationship was not strictly linear across the full signal range (**Fig. 4D**, top left). In *red1-mCIM*, desert bins showed a steeper dependence of S1-seq signal on Red1 ChIP, whereas island bins retained a shallower relationship similar to wild type (**Fig. 4D**, bottom left). When bins were grouped by Red1 ChIP level, the strongest dependence of S1-seq on Red1 was observed in low-Red1 regions, while middle- and high-Red1 regions showed weaker slopes (**Fig. 4D**, right). This suggests that DSB formation is particularly sensitive to Red1 abundance when Red1 levels are limiting, a condition enriched in desert regions after disruption of the Red1-CIM.

Together, these analyses indicate that the *red1-mCIM* mutation reduces DSB formation in a manner that broadly tracks local loss of Red1 chromatin association, with the strongest impact in Rec8-dependent domains. Thus, Red1-CIM-mediated interaction with Rec8 promotes DSB formation primarily by supporting Red1 recruitment to chromosomal regions where Red1 is limiting, while Rec8-independent mechanisms preserve Red1 binding and DSB formation at islands (see Discussion).

### Red1-CIM disruption causes severe chromosome segregation defects despite partial retention of DSB formation

To further evaluate the functional consequences of disrupting the Red1-CIM, we examined spore viability and chromosome segregation. Although both *red1-mCIM* and *red1-ΔCIM* mutants progressed through meiosis with kinetics comparable to wild type (**Supplemental Fig. S2A**), spore viability was strongly reduced in both mutants (**Fig. 5A**). Tetrad analysis revealed an overrepresentation of tetrads containing two or four dead spores, a pattern characteristic of chromosome missegregation during meiosis I (MI nondisjunction) (**Fig. 5A**).

**Figure 5.**
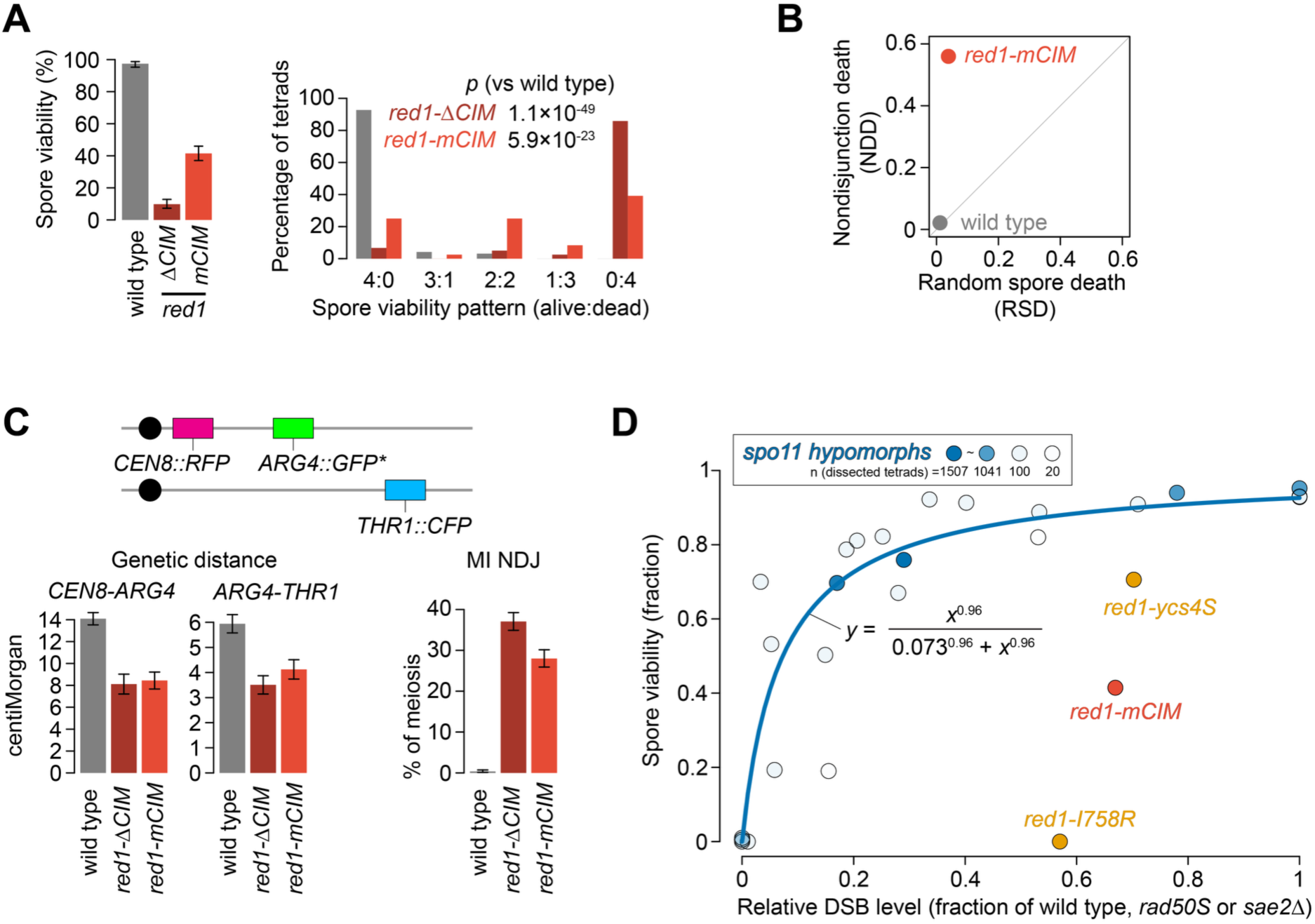
Red1-CIM mutants show severe post-DSB defects. **A.** Spore viability and tetrad viability patterns for wild type, *red1-ΔCIM*, and *red1-mCIM*. Error bars indicate 95% confidence intervals calculated assuming a binomial distribution. Fisher’s exact test was used to test differences in spore viability patterns relative to wild type (**Supplemental Table S1**). **B.** Analysis of spore death patterns using TetFit, separating meiosis I nondisjunction death (NDD) from random spore death (RSD). The *red1-ΔCIM* mutant was excluded from this analysis because its severe spore lethality falls outside the reliable range of the TetFit model. **C.** Schematic of the chr8 spore-autonomous fluorescence assay and quantification of genetic distances and meiosis I nondisjunction (MI-NDJ) (**Supplemental Table S2**). Error bars for genetic distance indicate standard error; error bars for MI-NDJ indicate 95% confidence intervals calculated assuming a binomial distribution. **D.** Relationship between DSB levels and spore viability. Spore viability is plotted as a function of relative DSB levels, expressed as a fraction of wild type, measured in *rad50S* or *sae2Δ* backgrounds as indicated (**Supplemental Table S3**). Data from *spo11* hypomorphic mutants are shown with a fitted Hill curve: SV = DSB^h / (K^h + DSB^h); K = 0.073, h = 0.96. Hill fitting was performed by weighted nonlinear regression, with weights proportional to the number of dissected tetrads contributing to each data point. Data points for *red1* mutants are plotted for comparison and were not included in the fit.

To quantify the contribution of MI nondisjunction to spore death, we applied TetFit (Chu and Burgess 2016). In *red1-mCIM* mutants, the majority of spore death was attributable to MI nondisjunction (∼56%) rather than random spore death (∼4%) (**Fig. 5B**). These results suggest that disruption of the Red1-CIM compromises meiotic chromosome segregation, most likely through defects in interhomolog recombination.

This conclusion was further supported by a spore-autonomous fluorescence assay on chr8 (Thacker et al. 2011), which revealed reduced genetic distance across both the *CEN8–ARG4* and *ARG4–THR1* intervals (∼60–70% of wild-type levels), together with a strong increase in MI nondisjunction (30–35%, corresponding to ∼80–100-fold over wild type) (**Fig. 5C**). Notably, this reporter interval overlaps an island-containing region, where Red1 ChIP and S1-seq signals were reduced but still partially retained in *red1-mCIM* (**Supplemental Fig. S5B**). Thus, the recombination and segregation defects observed at this interval occur in a region where Red1 binding and DSB formation are not completely abolished.

Together, these observations indicate that Red1-CIM disruption causes severe chromosome segregation defects, despite partial retention of Red1 binding and DSB formation at the chr8 reporter interval.

### The spore viability defect in *red1-mCIM* cannot be explained by reduced DSB formation alone

We next asked whether the spore viability defect in *red1-mCIM* could be explained solely by reduced DSB formation, or whether an additional defect acting after DSB formation contributes to spore death. To address this, we modeled the relationship between DSB levels and spore viability using published data from *spo11* hypomorphic mutants (Diaz et al. 2002; Martini et al. 2006; Rockmill et al. 2013) (**Fig. 5D**; **Supplemental Table S3**). Across these datasets, spore viability scaled with DSB levels and was well described by a Hill function (SV = DSB^h / (K^h + DSB^h); K = 0.073, h = 0.96). Because these datasets were collected in different genetic backgrounds and assays (*rad50S/sae2Δ*, Southern blotting vs S1-seq), we treat the fitted curve as an empirical reference rather than an absolute calibration.

This model indicates that substantial reductions in DSB formation can be tolerated with relatively modest effects on spore viability. For example, DSB levels around 50–70% of wild type are expected to support high (∼85–90%) spore viability in *spo11* hypomorphic backgrounds (**Fig. 5D**). In contrast, *red1-mCIM* exhibited only ∼41% spore viability despite retaining ∼67% of wild-type genome-wide DSB levels at 6 h in meiosis (**Fig. 3B**; **Fig. 5D**), representing a large deviation from the spore viability predicted by the empirical reference model (89%) (**Fig. 5D**).

We also plotted published data from additional *red1* alleles (Lin et al. 2010; Markowitz et al. 2017; Vale-Silva et al. 2019), which similarly exhibited lower spore viability than predicted based on their DSB levels alone (**Fig. 5D**; see Discussion). This suggests that the discrepancy between DSB levels and spore viability is not unique to *red1-mCIM*, but may reflect a broader role for Red1 in promoting productive DSB repair and crossover formation after DSBs are made.

Together, these observations indicate that the severe spore viability defect in *red1-mCIM* cannot be explained by reduced DSB formation alone and instead reflects additional defects acting downstream of DSB formation.

### The *red1-CIM* mutants fail to establish normal Mek1-dependent post-DSB signaling

A key post-DSB function of Red1 is activation of the meiosis-specific kinase Mek1, which promotes interhomolog recombination and enforces meiotic checkpoint signaling (Hollingsworth and Gaglione 2019). Mek1 activity is required for the characteristic prophase arrest observed in *dmc1Δ* cells and can be monitored by phosphorylation of known Mek1 substrates, including threonine 11 of histone H3 (H3-T11ph) (Kniewel et al. 2017).

To determine whether Red1-CIM mutations affect Mek1 activity, we examined H3-T11 phosphorylation during meiotic prophase. While H3-T11ph was readily detected in wild-type cells, both *red1-mCIM* and *red1-ΔCIM* mutants showed strongly reduced H3-T11ph levels across the time course examined (**Fig. 6A**). Because *red1-mCIM* retains substantial DSB formation, the loss of H3-T11ph in this mutant indicates that Red1-CIM disruption impairs post-DSB signaling in addition to reducing DSB formation.

**Figure 6.**
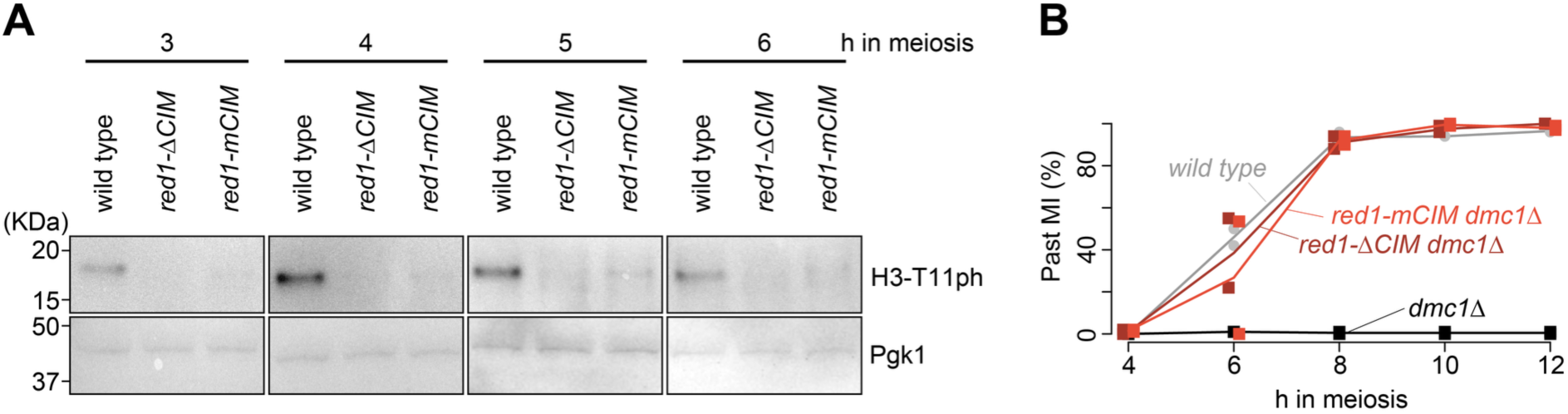
Red1-CIM is required for Mek1 activation during meiotic prophase. **A.** Immunoblot analysis of histone H3-T11 phosphorylation (H3-T11ph) in wild type, *red1-ΔCIM*, and *red1-mCIM* at the indicated times in meiosis. Pgk1 is a loading control. **B.** Meiotic progression in *dmc1Δ* backgrounds with or without Red1-CIM mutations, showing loss of the prophase arrest characteristic of Mek1 activation.

Consistent with this impaired signaling state, *red1-CIM* mutants also failed to maintain the meiotic prophase arrest characteristic of *dmc1Δ* cells. Whereas *dmc1Δ* cells exhibited a robust arrest, as expected for Mek1-proficient cells, both *red1-mCIM dmc1Δ* and *red1-ΔCIM dmc1Δ* strains progressed through meiosis with kinetics similar to wild type (**Fig. 6B**). Thus, Red1-CIM disruption compromises both Mek1 substrate phosphorylation and Mek1-dependent checkpoint maintenance.

Together, these findings indicate that the Red1-CIM is required to establish a normal Mek1-dependent post-DSB signaling state. Thus, in addition to promoting Red1 recruitment and DSB formation in Rec8-dependent chromosomal domains, the Red1–Rec8 interaction is important for post-DSB meiotic functions required for interhomolog recombination and faithful chromosome segregation (see Discussion).

### Conservation of Red1–Rec8 interaction in fungi

To address whether the Red1–Rec8 interaction is conserved beyond *S. cerevisiae*, we analyzed Red1-family proteins across nested fungal taxonomic ranks. This analysis showed that the CIM is conserved at all taxonomic levels from the Saccharomycetaceae family to the Fungi kingdom (**Fig. 7A,B**; **Supplemental Fig. S6**).

**Figure 7.**
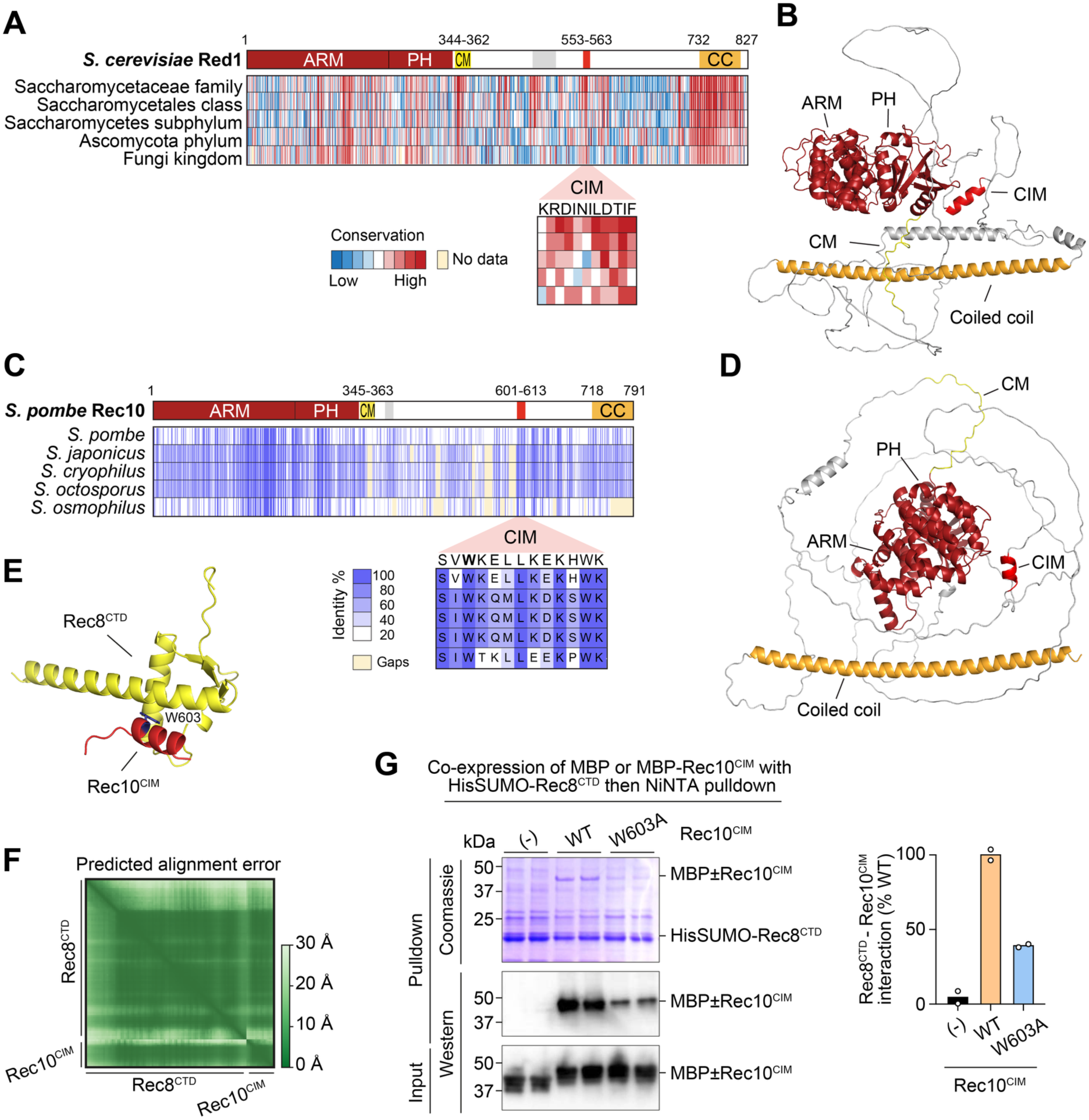
Conservation of Red1/Rec10-Rec8 interaction in *S. pombe*. **A.** Domain structure of *S. cerevisiae* Red1 and conservation analysis in fungi. **B.** AlphaFold model of Red1 (AF-P14291-F1-v6). **C.** Domain structure of *S. pombe* Rec10 and conservation analysis in *Schizosaccharomyces*. **D.** AlphaFold model of Rec10 (AF-Q09823-F1-v6). **E.** AlphaFold model of the Rec8 CTD (residues 469-561) bound to Rec10-CIM (residues 601-613). **F.** Predicted alignment error plot of the model shown in panel E. **G.** Pulldown analysis of MBP-tagged Rec10-CIM and HisSUMO-tagged Rec8-CTD. Plot shows the quantification of anti-MBP Western blot (mean from two replicates per construct (circles)).

Surprisingly, conservation of the Red1-CIM was lowest at the Ascomycota phylum level. This reduced signal was largely attributable to the composition of our Ascomycota dataset, in which approximately 60% of Red1 homologs belong to the Pezizomycotina subphylum.

Whereas a *S. cerevisiae*-type CIM was readily detected in Saccharomycotina, Pezizomycotina Red1 homologs lacked a clearly conserved CIM-like motif at the equivalent position. To examine this divergence further, we analyzed Red1 domain organization and sequence conservation in Pezizomycotina, using *Aspergillus fumigatus* Red1 as a representative example. This analysis revealed that Pezizomycotina Red1 proteins contain a markedly expanded intrinsically disordered region compared with Saccharomycotina homologs, with several conserved patches distributed across the IDR (**Supplemental Fig. S7**).

Notably, none of these conserved Pezizomycotina IDR patches aligned with the Saccharomycotina CIM in our cross-subphylum alignment, consistent with extensive sequence divergence and the approximately 2.7-fold expansion of the Pezizomycotina IDR. These observations suggest that Pezizomycotina Red1 proteins either harbor a highly diverged CIM that is not detectable from primary-sequence conservation alone, or have evolved a distinct mechanism for coupling Red1-family axis proteins to cohesin. Within the Taphrinomycotina subphylum, which includes the fission yeast *Schizosaccharomyces pombe*, clear Rec10 orthologs were restricted to *Schizosaccharomyces* species in our dataset, consistent with the deep divergence of fission yeast Rec10 from budding yeast Red1.

We therefore examined *S. pombe* Rec10 as a stringent test of conservation. Rec10 contains a predicted α-helix within its central IDR (residues 601-613) that is conserved across *Schizosaccharomyces* species (**Fig. 7C, D**, **Supplemental Fig. S8**). AlphaFold3 modeling placed this helix against the C-terminal domain of *S. pombe* Rec8 with high confidence (pTM 0.78, ipTM 0.79), in a configuration analogous to that predicted for the *S. cerevisiae* Red1–Rec8 complex (**Fig. 7E,F**). We therefore refer to this Rec10 segment as a putative CIM.

We tested the predicted interaction biochemically by co-expressing MBP-tagged Rec10-CIM with HisSUMO-tagged Rec8-CTD (residues 469-561) in *E. coli* and isolating HisSUMO-Rec8 by Ni-NTA pulldown. Rec10-CIM was efficiently recovered with Rec8-CTD, whereas alanine substitution of the conserved interface residue Rec10-W603 strongly reduced binding (**Fig. 7G**). Thus, Rec10-CIM is sufficient for direct interaction with the Rec8-CTD *in vitro*.

Together, these data identify a Rec8-binding CIM in *S. pombe* Rec10. Direct interaction between a Red1-family axis protein and the Rec8 CTD is therefore conserved between distantly related budding and fission yeasts, despite extensive divergence in primary sequence.

### Conservation of Red1–Rec8 interaction in plants

We next asked whether an analogous Rec8–axis interaction is present in plants. Plant Red1-family proteins, represented by *Arabidopsis thaliana* ASY3, preserve the long IDR and C-terminal coiled-coil architecture characteristic of meiotic axis proteins, although flowering plants and conifers have lost the N-terminal ARM and PH domains present in yeast Red1 (**Fig. 8A**). Within the ASY3 IDR, we identified a single predicted α-helix (residues 190–210) that is strongly conserved across plant taxonomic ranks, together with additional deeply conserved sequence blocks of unknown function (**Fig. 8A,B**; **Supplemental Fig. S9**).

**Figure 8.**
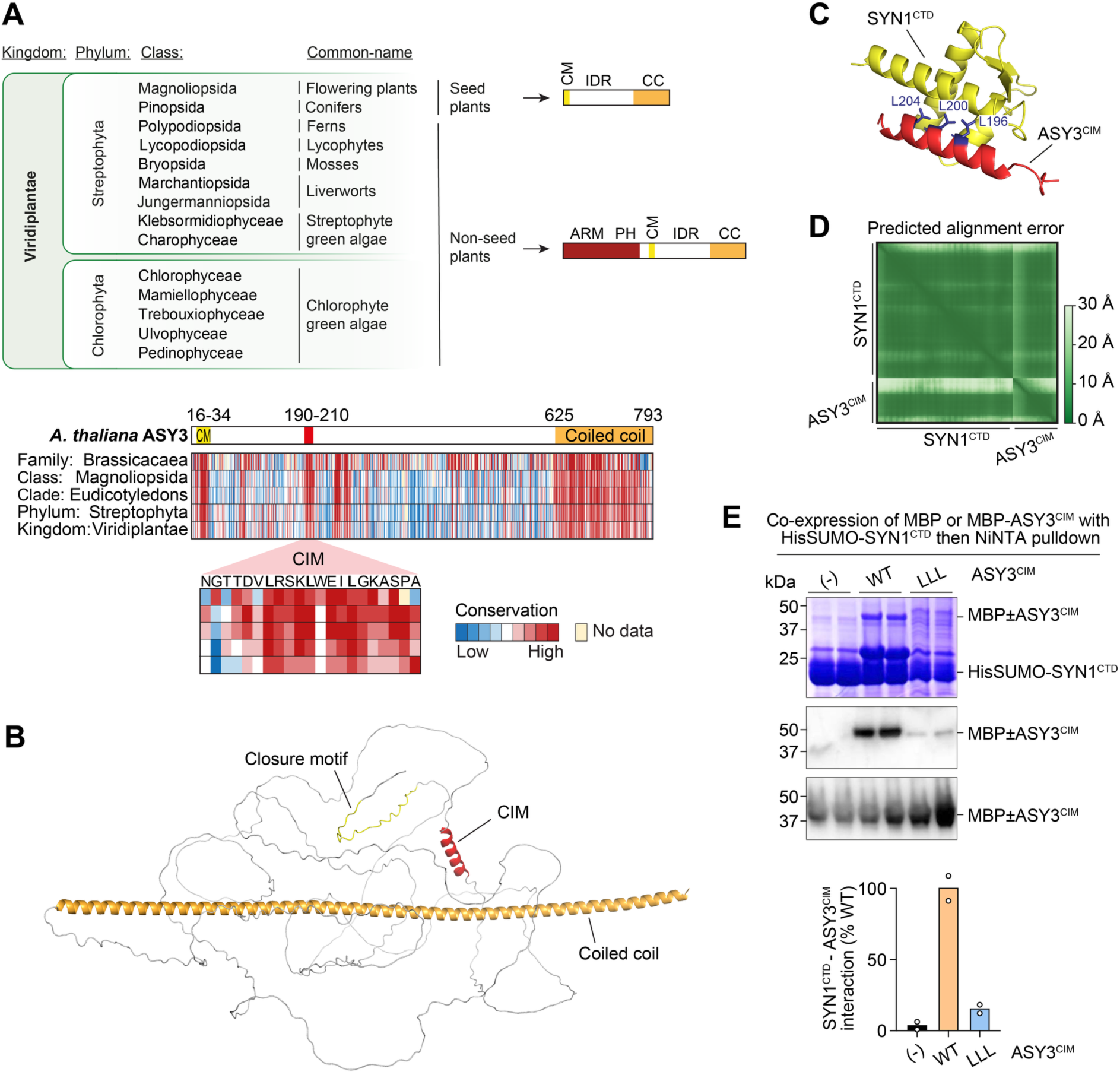
Conservation of Red1-Rec8 interaction in plants. **A.** Domain structure of Red1 orthologs in plants and conservation analysis of *A. thaliana* ASY3. **B.** AlphaFold model of ASY3 (AF-Q0WR66-F1-v6). **C.** AlphaFold model of the SYN1 CTD (residues 539-617) bound to ASY3-CIM (residues 190-210). **D.** Predicted alignment error plot of the model shown in panel C. **E.** Pulldown analysis of MBP-tagged ASY3-CIM and HisSUMO-tagged SYN1-CTD. Plot shows the quantification of anti-MBP Western blot (mean from two replicates per construct (circles)).

AlphaFold3 modeling predicted that the ASY3 helix docks onto the C-terminal domain of the Rec8 ortholog SYN1 in a configuration analogous to the fungal Red1/Rec10–Rec8 complexes (**Fig. 8A,C,D**). In the model, three deeply conserved leucine residues (L196, L200, and L204) in ASY3 form the hydrophobic core of the predicted interface with SYN1-CTD. We therefore refer to this ASY3 segment as a putative CIM.

Consistent with this model, MBP-tagged ASY3-CIM interacted with HisSUMO-tagged SYN1-CTD in pulldown assays, whereas alanine substitution of the three conserved leucines (LLL) markedly reduced binding (**Fig. 8E**). These data support conservation of a direct Red1/ASY3–Rec8/SYN1 interaction in plants, extending the Rec8–axis coupling mechanism beyond fungi.

Taken together with the fungal data, these results indicate that the direct coupling between Red1-family axis proteins and the Rec8-family kleisin CTD is broadly conserved, while allowing lineage-specific variation in the molecular details of the interface. This conserved Rec8–axis interaction likely represents a general mechanism by which cohesin is coupled to meiotic chromosome axis organization and recombination control.

## Discussion

### Red1-CIM defines a direct cohesin–axis molecular bridge

Meiotic chromosome axes provide a platform that coordinates DSB formation, interhomolog recombination bias, checkpoint signaling, and faithful chromosome segregation. In budding yeast, Rec8-containing cohesin has long been known to promote recruitment of the axis proteins Red1 and Hop1, but the molecular basis of this connection has remained unclear. Here, we identify a conserved motif in Red1, which we term the cohesin-interacting motif (CIM), that directly binds the C-terminal domain of Rec8. This interaction provides a molecular explanation for the Rec8-dependent pathway of Red1 recruitment to meiotic chromosomes.

The predicted Red1-CIM-binding surface on Rec8 is distinct from the Smc1-binding interface, suggesting that Red1 binding is compatible with the core architecture of the cohesin ring. This provides a simple mechanism by which Rec8 can function not only as a kleisin subunit mediating sister chromatid cohesion, but also as a meiosis-specific platform for axis protein recruitment. The absence of a comparable predicted interaction with Scc1 further suggests that this interface contributes to the meiosis-specific ability of Rec8-containing cohesin to support Red1 recruitment, consistent with previous observations that replacing Rec8 with Scc1 preserves cohesion but fails to restore normal Red1/Hop1 chromosome association (Brar et al. 2009; Sun et al. 2015).

### Red1 recruitment reflects variable contributions from Rec8-dependent and Rec8-independent pathways

Throughout this study, we used the island/desert classification as an operational framework to distinguish regions that retain relatively high Red1 binding when Rec8-dependent recruitment is impaired from regions that depend more strongly on Rec8 (**Fig. 2B**) (Heldrich et al. 2022). However, this classification should not be interpreted as a strict binary division. Red1 ChIP signal in *pREC8-SCC1* does not form two clearly separated populations, but instead shows a major low-signal population with a continuous high-signal tail (**Fig. 2F**), suggesting that chromosomal regions differ in the extent to which they support Rec8-independent Red1 recruitment. In addition, regions classified as islands are not necessarily independent of Rec8 in wild-type cells. Red1 profiles in wild type and *pREC8-SCC1* differ even within island regions (**Fig. 2C,F**; **Supplemental Fig. S3B**). Both *pREC8-SCC1* and *red1-mCIM* reduce Red1 binding in some island regions (**Fig. 2C,G**; **Supplemental Fig. S3B,C**). Islands likely receive Red1 through both Rec8-independent and Rec8-dependent pathways. Thus, islands are best viewed as regions with strong Rec8-independent recruitment superimposed on the normal Rec8-dependent pathway, rather than as regions where Rec8 makes no contribution.

Furthermore, loss of sharp Red1 peaks in *pREC8-SCC1* and *red1-CIM* mutants does not necessarily indicate complete loss of recruitment. In wild type, many peaks reflect TTS-proximal enrichment generated by transcription-dependent repositioning of cohesin and associated Red1 (**Fig. 2D,E**) (Sun et al. 2015). When this focusing is impaired, profiles become flatter, especially in deserts, yet desert signals in the mutant remain above the untagged control (**Fig. 2D**). Thus, the loss of distinct peaks can reflect reduced and spatially defocused Red1 binding rather than absence of Red1 (**Fig. 2C**).

Together, these observations suggest that Rec8-dependent and Rec8-independent pathways contribute in different proportions across chromosomal regions, creating a continuous spectrum of Red1 binding states. We therefore use the island/desert framework as an operational tool to highlight the role of the Red1-CIM in Rec8-dependent recruitment, while emphasizing that Red1 recruitment itself varies continuously across chromosomal regions.

### Red1–Rec8 interaction promotes DSB formation in Red1-limiting domains

Our S1-seq analysis indicates that the Red1-CIM is not absolutely required for Spo11-dependent DSB formation, consistent with previous work showing that DSBs still form in *red1Δ* and *rec8Δ* mutants, although their levels and distributions are altered (Klein et al. 1999; Kugou et al. 2009; Kim et al. 2010). Thus, the Red1–Rec8 interaction is unlikely to license DSB formation itself, but instead contributes to where DSBs form most efficiently.

Total V5-Red1 ChIP signal was reduced to ∼57% of wild type, whereas hotspot-associated S1-seq signal was retained at ∼86% of wild type at 3.5 h and ∼67% at 6 h (**Fig. 2A**; **Fig. 3B**). The larger reduction at 6 h may reflect premature loss of DSB competence, potentially linked to impaired post-DSB signaling (**Fig. 6**). Nevertheless, this non-proportional relationship is consistent with previous Red1 dosage studies, suggesting that relatively low levels of chromatin-associated Red1 can still support substantial DSB formation (Markowitz et al. 2017; Vale-Silva et al. 2019).

However, DSB reduction was not uniform. In *red1-mCIM*, local decreases in S1-seq signal broadly followed local decreases in Red1 ChIP signal, especially in Rec8-dependent deserts (**Fig. 4C**; **Supplemental Fig. S5A**). The Red1-ChIP–S1-seq relationship was nonlinear: DSB formation was most sensitive to Red1 levels in low-Red1 regions, whereas higher-Red1 regions showed a shallower dependence (**Fig. 4D**). Thus, islands, where Rec8-independent mechanisms maintain Red1, are relatively buffered, whereas deserts are more sensitive to disruption of the Red1–Rec8 interaction.

Together, these findings support a model in which the Red1-CIM promotes DSB formation in a quantitative and region-specific manner by maintaining sufficient Red1 in Rec8-dependent domains.

### Red1-CIM supports a post-DSB state required for Mek1 activation

Consistent with a post-DSB defect beyond reduced DSB formation (**Fig. 5**), *red1-CIM* mutants showed impaired Mek1 activation, as indicated by reduced H3-T11 phosphorylation and loss of the *dmc1Δ*-dependent prophase arrest (**Fig. 6**). Because Hop1 phosphorylation—a prerequisite for Mek1 recruitment and activation (Niu et al. 2005; Carballo et al. 2008)—can still occur in *spo11* hypomorphic backgrounds with reduced DSB levels (Lin et al. 2010), the Mek1 defect in *red1-mCIM* is unlikely to be explained simply by reduced DSB formation.

This defect is also unlikely to be explained by loss of the CIM–Rec8 interaction alone. Although Rec8 shapes Red1 distribution (Klein et al. 1999; Sun et al. 2015), Mek1-dependent signaling can still be detected in *rec8Δ* cells: Hed1-T40 phosphorylation is retained in *rec8Δ* cells (Callender et al. 2016), and Mek1-dependent recombination control can occur in genetic contexts lacking Rec8 cohesin (Callender and Hollingsworth 2010; Kim et al. 2010). Thus, the Mek1 defect in *red1-CIM* mutants likely reflects a role of the CIM beyond Rec8-dependent Red1 positioning, contributing to a signaling-competent Red1 state.

One possibility is that *red1-mCIM* reduces chromatin-associated Red1 below the level required for efficient signaling. This would be consistent with *red1-ycs4S*, in which reduced Red1 dosage disproportionately impairs meiotic checkpoint arrest relative to DSB formation (Markowitz et al. 2017; Vale-Silva et al. 2019). However, *red1-mCIM* retains more chromatin-associated Red1 than *red1-ycs4S* and preserves substantial DSB formation (**Fig. 2A**; **Fig. 3B**), suggesting that reduced Red1 abundance alone is unlikely to explain the Mek1 defect.

An alternative, non-mutually exclusive possibility is that the CIM contributes to a qualitative Red1 state required for post-DSB signaling. Several Red1 alleles support the idea that Mek1 activation requires more than Red1 chromosomal association. The *red1-I758R* mutant disrupts SUMO-chain binding, impairs Red1 filament-bundle formation, and abolishes Hop1 phosphorylation despite allowing recombination initiation (Lin et al. 2010; Cheng et al. 2013; West et al. 2019). Similarly, *red1-I743A* disrupts Ddc1 interaction and impairs Red1 filament formation, linking the Red1 C-terminal coiled-coil region to both 9-1-1-dependent checkpoint signaling and higher-order Red1 organization (Eichinger and Jentsch 2010; Cheng et al. 2013; West et al. 2019). Thus, Red1 abundance, SUMO-dependent interactions, 9-1-1 association, filament formation, and CIM-dependent interactions may together contribute to a signaling-competent Red1 state.

Together, these findings suggest that the Red1-CIM is required not only for efficient Red1 recruitment and DSB patterning, but also for establishing a post-DSB Red1 state that supports Mek1 activation. In this view, *red1-mCIM* cells make substantial numbers of DSBs, but these DSBs fail to fully engage the Red1-dependent signaling environment required for normal interhomolog recombination outcomes.

### Limitations and allele-specific considerations

Several caveats are important for interpreting these results. First, although both *red1-mCIM* and *red1-ΔCIM* disrupt the CIM region, the two alleles are not equivalent. The *red1-ΔCIM* mutant showed reduced nuclear localization of Red1, likely because deletion of the CIM region also disrupts a putative nuclear localization signal. Therefore, we place greater mechanistic weight on *red1-mCIM*, which preserves Red1 protein abundance and nuclear enrichment while disrupting the conserved interface residues. The broader defects observed in *red1-ΔCIM* likely reflect a combination of impaired Rec8 interaction and reduced nuclear Red1 availability.

Second, although we generated reciprocal Rec8-side interface mutants, these alleles reduced Rec8 protein levels (**Supplemental Fig. S2F**) and were therefore difficult to interpret. Because Rec8 is a core cohesin subunit, changes in its abundance could affect cohesin dosage or stability independently of Red1 binding. We therefore focused on the Red1-side allele *red1-mCIM* as the more interpretable *in vivo* perturbation for dissecting the Red1–Rec8 interaction.

Third, ChIP-seq measures chromatin-associated Red1 abundance, not the physical architecture of the meiotic axis directly. Therefore, when we refer to Red1 chromosomal states, we do so operationally, based on Red1 abundance, Rec8 dependence, DSB output, and Mek1 signaling behavior. Thus, our ChIP-based analyses define functional Red1 chromosomal states, but do not directly resolve higher-order axis architecture.

### Evolutionary conservation and broader implications

The evolutionary analyses extend the significance of the Red1-CIM beyond budding yeast. In *S. cerevisiae*, the CIM provides a direct molecular bridge between Rec8-containing cohesin and the Red1-Hop1 chromosome axis. The identification of analogous Rec8-binding helical motifs in *S. pombe* Rec10 and *Arabidopsis thaliana* ASY3 suggests that direct cohesin–axis coupling is a recurrent and likely widespread strategy for building meiotic chromosome axes in fungi and plants. Importantly, the conserved feature is not a strict amino-acid sequence, but a short helical module embedded in the IDR of axis proteins that docks onto the Rec8-family kleisin CTD.

This model has two broader implications. First, Rec8-family kleisins should be viewed not only as meiosis-specific cohesion subunits, but also as molecular adaptors that help position axis proteins on chromosomes. Through this adaptor function, Rec8 can influence both where DSBs form and how DSB-associated signaling is executed. Second, the divergence observed among fungi and plants suggests that meiotic chromosome axes can preserve a common architectural principle while diversifying the molecular details of the cohesin–axis interface.

In budding yeast, the CIM is essential for efficient Red1 recruitment in Rec8-dependent domains, for the spatial distribution of DSB formation, and for establishing the post-DSB signaling state required for Mek1 activation, interhomolog recombination, and accurate chromosome segregation. Future *in vivo* analyses of the corresponding CIMs in plants will determine how broadly this cohesin-axis mechanism controls recombination patterning and chromosome segregation beyond fungi.

### Materials and Methods

#### Yeast strains and plasmids

All *S. cerevisiae* and *S. mikatae* strains and plasmids used in this study are listed in **Supplemental Tables S4 and S5**. *S. cerevisiae* strains were derived from the SK1 background (Kane and Roth 1974), and *S. mikatae* strains from IFO1815 (Scannell et al. 2011).

Gene deletions and point mutations were introduced by standard PCR-based gene targeting (Longtine et al. 1998) using a lithium acetate transformation method (Gietz and Schiestl 2007). Transformants were verified by PCR and Sanger sequencing. The *pREC8-SCC1-3HA* strain (Toth et al. 2000) was provided by the Adele Marston laboratory. The *S. mikatae* spike-in control strains were generated by replacing the *RAD50* stop codon and the *SAE2* open reading frame with V5-KanMX and KanMX, respectively.

To generate *red1* mutant plasmids, the N-terminally V5-tagged *RED1* coding sequence was amplified from the *V5-RED1* strain provided by the Franz Klein laboratory (Lin et al. 2010; Sun et al. 2015) and cloned into a pUG27-derived vector carrying a loxP-flanked *KanMX* cassette (Gueldener et al. 2002). The *RED1* promoter (677 bp upstream of the start codon) and terminator (497 bp downstream of the stop codon) regions were then inserted to yield pHM27, carrying *pRED1-V5-RED1-loxP-KanMX-loxP-tRED1* (**Supplemental Table S5**).

The *red1-ΔCIM* allele, deleting residues 553–563, and the *red1-mCIM allele*, carrying seven substitutions, were introduced by site-directed mutagenesis (Ho et al. 1989) to yield pHM28 and pHM29, respectively (**Supplemental Table S5**).

To generate V5-tagged and untagged wild-type and mutant strains, restriction enzyme-digested plasmids were used to replace the endogenous *RED1* locus. G418-resistant transformants were verified by PCR and Sanger sequencing.

The *rec8* mutant plasmids were generated by site-directed mutagenesis (Ho et al. 1989) to introduce the *F581A*, *Y584A*, and *P613K* substitutions into a wild-type *REC8* plasmid carrying the *REC8* promoter (862 bp upstream of the start codon, SK1), coding sequence (SK1), and terminator (884 bp downstream of the stop codon, W303) with the *KlTRP1* marker inserted downstream of the stop codon. This yielded pHM75, pHM76, and pHM64, respectively (**Supplemental Table S5**). The *rec8-CIM* plasmid (pHM65) encodes full-length Rec8 fused at its C-terminus, via a flexible linker (GGGGGS), to Red1 residues 543–564 containing the CIM. To generate *rec8* mutant yeast strains, plasmids were cleaved within the promoter and terminator regions with the indicated restriction enzymes (**Supplemental Table S5**), and the resulting fragments were used to transform a *rec8Δ* strain generated by replacing the *REC8* open reading frame with *KanMX6*.

#### Sporulation and meiotic time courses

Synchronous meiotic cultures of *S. cerevisiae* and *S. mikatae* were prepared using the SPS pre-growth method as described previously (Mu et al. 2020; Murakami et al. 2020). Briefly, diploid cells were selected for respiratory competence on YPG plates, grown on YPD plates, and then cultured sequentially in YPD and SPS medium. Cells were transferred to SPM at a density of 4×10^7^ cells/mL to induce meiosis; the time of transfer to SPM was defined as t = 0 h. Cultures were incubated at 30°C with shaking, and samples were collected at the indicated time points.

To assess meiotic progression, aliquots were fixed in 50% ethanol and stained with DAPI. Mono-, bi-, and tetranucleate cells were scored by fluorescence microscopy. More than 100 cells were scored per strain per time point for each of two biological replicates.

#### Immunoblotting

Whole-cell extracts were prepared by TCA extraction as described previously (Kniewel et al. 2017), with minor modifications. Briefly, 3 mL of meiotic culture was harvested at the indicated time points, fixed in 20% TCA, disrupted by bead beating, and protein pellets were resuspended in Laemmli sample buffer.

Proteins were separated on NuPAGE Bis-Tris gels (Invitrogen) and transferred to PVDF membranes (Cytiva, 10600022). V5-Red1 and Rec8 were resolved on 4–12% gradient gels, whereas H3-T11ph was resolved on 12% gels. Membranes were blocked in 5% non-fat milk/TBST and incubated with primary antibodies against V5 (Bio-Rad, MCA1360; 1:1,000), H3-T11ph (EMD Millipore, 04-789; 1:1,000), or Pgk1 (Invitrogen, PA5-28612; 1:2,000 or 459250; 1:10,000 for **Supplemental Fig. S2F**). Rec8 was detected using rabbit anti-Rec8 ((Bommi et al. 2019); 1:1,000). HRP-conjugated anti-rabbit (Invitrogen, 31460) and anti-mouse (Cytiva, NA931V) secondary antibodies were used at 1:2,000, and signals were detected using Immobilon Western Chemiluminescent HRP Substrate before imaging with a Syngene G system.

#### Immunofluorescence microscopy and Red1 nuclear localization analysis

Whole-cell immunofluorescence staining was performed as described previously (Henggeler et al. 2025), with minor modifications. Meiotic cells were harvested 3 h after meiotic induction and fixed with 1% formaldehyde for 30 min at room temperature (see V5-Red1 ChIP-seq below for detail). Spheroplasting was performed with 50 μg/mL Zymolyase 100T at 30°C for 5 min, a condition optimized by monitoring sarcosyl lysis. Spheroplasts were adhered to poly-L-lysine slides and stained overnight with mouse anti-V5 antibody (Bio-Rad, MCA1360; 1:200), followed by Alexa Fluor 488-conjugated goat anti-mouse IgG (Invitrogen, A-11001; 1:500). Slides were mounted in Fluoroshield Mounting Medium with DAPI (Abcam, AB104139) before imaging.

Images were acquired on a Zeiss LSM710 inverted confocal microscope using a 63×/1.4 NA Plan-Apochromat oil-immersion objective, with identical acquisition settings across samples. Maximum-intensity projections of z-stacks were used for downstream analysis, with DIC images acquired in parallel for cell boundary segmentation.

Red1 nuclear localization was quantified using a custom CellProfiler v4.2.8.1 pipeline developed in this study. Nuclei were segmented from DAPI images: speckle features were enhanced (EnhanceOrSuppressFeatures, Tubeness method, feature size 35 pixels) and primary objects in the 30–100 pixel diameter range were identified by global intensity thresholding with intensity-based declumping; objects touching the image border were discarded. Cell boundaries were segmented from DIC images using Cellpose v3 with the pretrained ’cyto’ model (flow threshold 0.8, minimum size 70 pixels), supplying the DAPI image as a nuclei reference, called via the RunCellpose CellProfiler plugin. Nuclei were related to their parent cell, and only cells containing exactly one nucleus, with the nucleus fully contained within the cell boundary (filtered by AreaShape FormFactor), were retained. Cytoplasmic regions were defined as the cell area excluding the nucleus (IdentifyTertiaryObjects). For each retained cell, mean Alexa Fluor 488 (V5-Red1) intensity was measured in the nuclear and cytoplasmic regions (MeasureObjectIntensity), and the nuclear/cytoplasmic mean intensity ratio was calculated per cell (CalculateMath). Per-cell ratios were log2-transformed for plotting.

#### V5-Red1 ChIP-seq

V5-Red1 ChIP-seq with *S. mikatae* spike-in calibration was performed as described previously (Murakami and Keeney 2014; Huang et al. 2025) with modifications detailed below.

For each ChIP sample, 15 mL of meiotic *S. cerevisiae* culture (∼6×10^8^ cells) was harvested at 3 h after meiotic induction and cross-linked with 1% formaldehyde for 30 min at room temperature with rotation at 15 rpm and a 160° tilt, followed by glycine quenching (131 mM, 5 min). Cells were washed once with 25 mL of ice-cold TBS and stored at –80°C. For spike-in calibration, *S. mikatae* cells expressing Rad50-V5 (HMY390) were sporulated in parallel using the same SPS pre-growth protocol, harvested at 3 h, and fixed under identical conditions; 6×10^7^ *S. mikatae* cells (10% of the *S. cerevisiae* cell number) were added to each sample prior to lysis.

Cell pellets were resuspended in 300 μL of ice-cold lysis buffer (50 mM HEPES-KOH pH 7.5, 140 mM NaCl, 1 mM EDTA, 1% Triton X-100, 0.1% sodium deoxycholate) supplemented with 1 mM PMSF, 7 μg/mL aprotinin, 1% protease inhibitor cocktail (Sigma, P8215), and 1× Roche Complete EDTA-free protease inhibitor cocktail. Cells were disrupted by bead beating with 0.5 mm zirconia/silica beads (∼500 μL; BioSpec, 11079105z) on a FastPrep-24 instrument (MP Biomedicals) at 6.5 m/s for two 1-min cycles, with samples chilled on ice for 5 min between cycles. Chromatin was sheared on a Bioruptor Pico 2 (Diagenode) in 1.5 mL Bioruptor Pico tubes (Easy Mode, 30 s ON / 30 s OFF, 14 cycles, 4°C) to yield an average DNA size of ∼150 bp. Cell debris was pelleted by centrifugation at 15,000 rpm for 5 min at 4°C. A 20 μL aliquot of the clarified supernatant was retained as the Input sample and mixed with 100 μL of TE / 1.2% SDS; the remaining supernatant was taken forward for immunoprecipitation.

For immunoprecipitation, 4 μg of mouse anti-V5 antibody (Bio-Rad, MCA1360) was pre-incubated with each lysate at 4°C for 5 h with rotation. In parallel, 25 μL of Dynabeads Protein G (Invitrogen, 100.04D) per sample was washed twice with 400 μL of PBS containing 5 mg/mL BSA (PBS/BSA) and blocked in PBS/BSA at 4°C for 5 h with rotation. The lysate-antibody mixture was then added to the blocked beads and incubated overnight at 4°C with rotation. Bead washes and elution were carried out as described in Murakami and Keeney (2014), and the eluate was supplemented with 80 μL of TE / 1% SDS. Cross-links were reversed and RNA was digested by overnight incubation at 65°C with 1 μL of RNase A (10 mg/mL). Protein was subsequently digested with 7.5 μL of 20 mg/mL proteinase K at 50°C for 2 h. DNA was purified using the Monarch PCR & DNA Cleanup Kit (NEB, T1030L) following the manufacturer’s protocol and eluted in 50 μL of TE buffer.

Purified ChIP and Input DNA fragment sizes were confirmed on an Agilent TapeStation 4150 using the High Sensitivity D5000 ScreenTape assay (Agilent, 5067-5592). Samples were then adjusted to 100 μL with TE buffer and subjected to a second round of sonication on a Bioruptor Pico 2 (Diagenode) under the same conditions as the first sonication, except that 8 cycles were used, yielding final fragment sizes of ∼150 bp suitable for library preparation. Sequencing libraries were prepared from 100 μL of fragmented DNA using the NEBNext Ultra II DNA Library Prep Kit for Illumina (NEB, E7645S) with NEBNext Sample Purification Beads (NEB, E7104S) and NEBNext Multiplex Oligos for Illumina (Dual Index Primers Set 1, NEB, E7600S) following the manufacturer’s instructions. Libraries were sequenced by Macrogen on an Illumina NovaSeq X platform with paired-end 150 bp reads.

#### ChIP-seq data processing and spike-in normalization

Paired-end sequencing reads were quality- and adapter-trimmed using fastp v1.1.0 (-q 20 -l 32 -3 -c -p) (Chen et al. 2018). Trimmed reads were first aligned with Bowtie2 v2.5.5 (Langmead and Salzberg 2012) to a combined reference containing the *S. cerevisiae* SK1 PacBio assembly (GenBank GCA_002057885.1) and the *S. mikatae* IFO1815 assembly (available from the *Saccharomyces sensu stricto* resource at https://sss.genetics.wisc.edu). Alignment was performed using stringent perfect-match settings (--very-sensitive -L 32 -N 0 --score-min L,0,0 --no-discordant --no-mixed --dovetail -I 0 -X 500) to assign each read pair unambiguously to its strain of origin. Read pairs aligning equally well to both genomes, identified by the XS:i:0 tag on either mate, were removed using a custom awk filter. The remaining alignments were retained as primary, properly paired alignments using SAMtools v1.23 (samtools view -b -f 2 -F 256) (Danecek et al. 2021).

Reads assigned to *S. mikatae* chromosomes were counted to derive per-sample spike-in normalization factors. Reads assigned to SK1 chromosomes were extracted, converted back to paired FASTQ, and remapped to the *S. cerevisiae* S288C/sacCer2 reference genome (UCSC) using Bowtie2 with SNP-tolerant settings (--very-sensitive -L 32 -N 1 --no-discordant --no-mixed --dovetail -I 0 -X 500, with the default --score-min L,-0.6,-0.6, allowing approximately 15 mismatches per 150-bp read) to accommodate sequence differences between SK1 and S288C. Uniquely mapped, primary, properly paired alignments were retained (samtools view -b -q 20 -f 2 -F 256), and per-base coverage tracks were generated using SAMtools.

Coverage tracks were calibrated by normalization to one million uniquely mapped *S. mikatae* spike-in reads. Calibrated ChIP tracks were then divided by the corresponding calibrated input tracks to generate spike-in-normalized ChIP coverage maps, which were used for downstream analyses in R version 4.3.2.

#### S1-seq

S1-seq was performed as described previously (Mimitou and Keeney 2018), with the modifications described below. All strains used for S1-seq, including the spike-in control, were *sae2Δ* to preserve unresected DSB ends. *S. mikatae* was used as an exogenous spike-in control at a 1:5 ratio (*S. mikatae*:*S. cerevisiae*) to enable normalization across samples; spike-in cells were harvested 6 h after meiosis induction for all experiments.

Two library preparation workflows were used depending on the time point collected. For the 6 h time point, 8×10^8^ *S. cerevisiae* cells were mixed with 1.6×10^8^ *S. mikatae* spike-in cells and co-embedded to produce 8–10 agarose plugs. All available plugs were used for library preparation. For the 3.5 h time point, 4×10^8^ *S. cerevisiae* cells were mixed with 8×10^7^ *S. mikatae* spike-in cells and co-embedded to produce 3–5 agarose plugs, of which only 3 were used for library preparation. This provided sufficient DNA input after size selection, which imposes a maximum DNA quantity constraint regardless of plug number.

The protocol of Mimitou and Keeney (2018) was modified as follows. Section 5.2.1, steps 1–3 and 14–15: GELase was replaced by β-Agarase I (NEB) at 2 U per plug (or 1 U per 100 mg agarose gel), and digestion was carried out at 42°C for 16–24 h. Section 5.2.1, steps 11–12: Ethidium bromide was replaced by Midori Green Advanced (GeneFlow) for nucleic acid staining, and gels were visualized using a blue-light emitter (400–490 nm). Section 5.2.1, steps 16–19: Phenol:chloroform extraction was replaced by sodium acetate/isopropanol precipitation. Briefly, 0.1 volume of 3 M sodium acetate (pH 5.5) was added to the DNA sample and chilled on ice for 15 min. Samples were centrifuged at 15,000×*g* for 15 min, after which 2 volumes of 100% isopropanol were added to the supernatant and incubated at −20°C for at least 2 h. The precipitate was collected by centrifugation at 15,000×*g* for 15 min and washed once with 70% isopropanol. Section 5.2.1, step 21: The Chroma Spin-1000 column step was omitted.

Section 5.2.2, step 23: The Covaris shearing step was replaced by shearing with a Bioruptor Pico (Diagenode) in Easy Mode for 15 cycles (30 s ON / 30 s OFF). Size selection (post-shearing, replacing section 5.2.2, step 23 size selection): Fragment size selection targeting 200–500 bp was performed using SPRIselect beads (Beckman Coulter) according to the manufacturer’s instructions. For the 6 h samples, 100 µL of DNA input was used with 50 µL of beads; for the 3.5 h samples, 200 µL input was used with 100 µL of beads to maximize DNA recovery.

Section 5.4.2, steps 1–2: Rather than splitting the 20 µL streptavidin bead suspension into two parallel PCR reactions, only one reaction was performed using 10 µL of bead suspension as template. Library amplification was carried out using PCRBIO VeriFI™ polymerase (PCR Biosystems) instead of Phusion, under the following conditions: 98°C for 2 min; 14 cycles of 98°C for 30 s, 58°C for 30 s, 72°C for 15 s; final extension at 72°C for 10 min. Section 5.4.2, steps 6–24: The post-PCR polyacrylamide gel electrophoresis, gel extraction, and ethanol precipitation steps were replaced by size selection using SPRIselect beads (Beckman Coulter) targeting fragments of 200–300 bp, according to the manufacturer’s instructions. Libraries were submitted for commercial next-generation sequencing on an Illumina NovaSeq X platform with 150 bp paired-end reads, targeting 20–40 million raw reads per sample.

#### S1-seq data processing

Read trimming, mapping to the combined *S. cerevisiae* SK1 and *S. mikatae* reference genomes, and remapping of *S. cerevisiae* reads to the sacCer2 reference genome were performed as described in the ChIP-seq data processing and spike-in normalization section.

For S1-seq, the 5′ end of Read 2 corresponds to the S1 nuclease cleavage site in this library design. Read 2 alignments were therefore extracted from sacCer2-mapped BAM files, excluding secondary alignments, and 5′-end coverage was calculated at single-nucleotide resolution separately for the forward and reverse strands using bedtools genomecov -5 -d - strand (bedtools v2.31.1) (Quinlan and Hall 2010).

We used R for the following data processing. Top- and bottom-strand maps were shifted to represent the inferred Spo11 cleavage positions and then merged to generate genome-wide S1-seq signal tracks. Analogous strand-separated 5′-end coverage tracks were generated for *S. mikatae* reads using the combined-reference mapping coordinates directly, without remapping. Enrichment of *S. mikatae* signal at published meiotic hotspots (Lam and Keeney 2015) was used to verify that the spike-in DNA contributed genuine S1-seq signal.

We also performed de novo hotspot calling on wild-type and *red1-mCIM* S1-seq datasets using an approach similar to that described previously (Pan et al. 2011). However, we did not identify reproducible mutant-specific hotspots outside the published Spo11-oligo hotspot set (Mohibullah and Keeney 2017). Therefore, downstream hotspot-based analyses used the fixed Spo11-oligo hotspot annotation described below.

#### S1-seq background subtraction and spike-in calibration

Background signal was estimated separately for *S. cerevisiae* and *S. mikatae* using genomic positions outside annotated Spo11-oligo hotspots. Hotspot intervals were defined using published *S. cerevisiae* Spo11-oligo hotspots (Mohibullah and Keeney 2017) and *S. mikatae* Spo11-oligo hotspots (Lam and Keeney 2015).

For each sample, the distribution of per-position S1-seq counts in non-hotspot regions was summarized using a frequency table. Background cleavage was modeled as a Poisson process, and the background rate λ was estimated as:

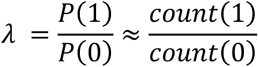

where count(1) and count(0) are the numbers of genomic positions with one and zero reads, respectively. The estimated λ was subtracted uniformly from all genomic positions in the corresponding dataset.

For spike-in calibration, background-subtracted *S. mikatae* signal was summed over annotated *S. mikatae* hotspot intervals. Hotspots with negative background-subtracted signal in any sample were excluded from the calibration-factor calculation. Sample-specific scaling factors were then calculated from the total background-subtracted *S. mikatae* hotspot signal. Background-subtracted *S. cerevisiae* signals were normalized to one million background-subtracted *S. mikatae* hotspot signal: calibrated signal = 10^6^×[background-subtracted *S. cerevisiae* signal] / [total background-subtracted *S. mikatae* hotspot signal].

For *S. cerevisiae* downstream analyses, hotspot-associated S1-seq signal was quantified using the fixed Spo11-oligo hotspot set from Mohibullah and Keeney (2017). This published annotation was used as a common reference set to enable direct comparisons across genotypes and time points. For each hotspot, hotspot-associated signal was calculated by summing calibrated S1-seq signal within ±100 bp of the smoothed Spo11-oligo peak. De novo hotspot calling on individual S1-seq datasets was not used for the primary analysis because peak detection was sensitive to overall signal strength and genotype-specific differences in DSB levels. For all downstream quantitative analyses, except for visualization of S1-seq profiles along chromosomal regions (**Fig. 3C**; **Fig. 4A**; **Supplemental Fig. S4C; Supplemental Fig. S5B**), we used hotspot-associated S1-seq signal.

#### Spore viability and tetrad analysis

Sporulation was performed as described above. After 24–48 h in SPM, sporulated cultures were treated with Zymolyase 100T in 1 M sorbitol at 30°C for 15 min, and tetrads were dissected on YPD plates using a Singer MSM 400 dissection microscope. Plates were incubated at 30°C for 2 days, and spore viability was scored by colony formation (**Supplemental Table S1**).

#### Spore-autonomous fluorescence reporter assay

Yeast cells were sporulated for two days and subjected to subsequent analysis to measure genetic distance and MI nondisjunction frequencies on chr8 as described previously (Thacker et al. 2011; Huang et al. 2025) and in **Supplemental Table S2**.

MI-NDJ values from the spore-autonomous fluorescence assay represent apparent chr8 MI nondisjunction frequencies. NPD events in the *ARG4–THR1* interval produce fluorescence segregation patterns indistinguishable from MI nondisjunction and were therefore included in the MI-NDJ class (Thacker et al. 2011). Consequently, these values may overestimate true chr8 MI nondisjunction. In *red1-mCIM,* the apparent MI-NDJ frequency measured by this assay (28.0%) was higher than the genome-average per-chromosome MI-NDJ estimate obtained from TetFit analysis (10.8%). This difference may reflect the chr8-specific nature of the fluorescence assay, the inclusion of *ARG4–THR1* NPD events in the apparent MI-NDJ class, and the model-based genome-average assumptions of TetFit.

#### DSB–spore viability modeling

The relationship between DSB levels and spore viability was modeled using published datasets from *spo11* hypomorphic mutants (Diaz et al. 2002; Martini et al. 2006; Rockmill et al. 2013; **Supplemental Table S3**). Relative DSB levels and spore viability values were converted to proportions by dividing percentage values by 100. DSB levels were normalized to the corresponding wild-type value within each study.

The *spo11* hypomorph data were fitted using a Hill function:

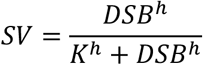

where *SV* is spore viability, *DSB* is the relative DSB level, *K* is the DSB level at half-maximal spore viability, and *h* is the Hill coefficient. Weighted nonlinear regression was performed in R using the nlsLM function from the minpack.lm package, with the number of dissected tetrads used as weights. The fitted parameters were *K* = 0.073 and *h* = 0.96.

#### Statistical analysis

Statistical analyses were performed in R (R version 4.3.2).

#### Expression plasmids

Plasmids are listed in **Supplemental Table S5**. Oligos are listed in **Supplemental Table S6**. Vectors for co-expression of Red1-CIM and Rec8-CTD orthologs were based on a pETDuet1 vector with the sequence coding a HisSUMO tag cloned into site 1 and an MBP tag cloned into site 2. Sequences coding for Rec8-CTD orthologs were codon-optimized for expression in *E. coli*, synthesized as gBlocks (IDT), and cloned downstream of the HisSUMO tag by Gibson assembly. Sequences of the synthetic DNA fragments are listed in **Supplemental Table S7**.

The plasmid for co-expression of *S. cerevisiae* HisSUMO-Rec8-CTD with MBP alone is pCCB1114. The Red1-CIM was then added downstream of the MBP coding sequence by inverse PCR of pCCB1114 with primers cb1627 and cb1628 followed by blunt-end ligation to yield pCCB1124. Mutant constructs were generated by inverse PCR and blunt-end ligation as follows (with resultant plasmids and PCR primers in brackets): Red1-F563A (pTR23, primers tr49 & tr50), Rec8-P613K (pTR24, primers tr32 & tr33), Rec8-I636A (pTR25, primers tr34 & tr35), Rec8-F581A (pTR26, primers tr37 & tr44), Rec8-Y584A (pTR27, primers tr51 & tr52), Rec8-Red1 fusion (pTR29, primers tr40 & tr41).

The plasmid for co-expression of *S. pombe* HisSUMO-Rec8-CTD with MBP alone is pCCB1116. The Rec10-CIM was then added downstream of the MBP coding sequence by inverse PCR of pCCB1116 with primers cb1629 and cb1630 followed by blunt-end ligation to yield pCCB1125. The Rec10-W603A mutation was generated by inverse PCR and blunt-end ligation using pCCB1125 as template and primers tr77 & tr78 to yield pTR42.

The plasmid for co-expression of *A. thaliana* HisSUMO-SYN1-CTD with MBP alone is pCCB1122. The ASY3-CIM was then added downstream of the MBP coding sequence by inverse PCR of pCCB1122 with primers cb1625 and cb1626 followed by blunt-end ligation to yield pCCB1123. The ASY3-L196A,L200A,L204A mutation was generated by inverse PCR and blunt end ligation using pCCB1122 as template and primers tr79 & tr80 to yield pTR43.

#### Pulldown assays

Plasmids to co-express HisSUMO-tagged Rec8-CTD and MBP-tagged Red1-CIM and their variants were transformed in *E. coli* BL21 DE3 cells. Transformants were selected using ampicillin and single colonies were transferred to flasks containing 50 ml liquid LB with ampicillin and grown at 37 °C. When the optical density at 600 nm (OD600) reached 0.6, 0.3 mM isopropyl β-D-1-thiogalactopyranoside (IPTG) was added, and cultures were transferred to a 16 °C shaker for overnight incubation (16–18 h). Cells were harvested and washed once with ice-cold 1X PBS, then centrifuged at 5000 rpm for 10 minutes at 4 °C. For pulldowns with *S. cerevisiae* proteins, the cell pellet was resuspended in ice-cold amylose buffer (25 mM HEPES-NaOH, pH 7.5, 500 mM NaCl, 10% glycerol, 1 mM DTT, 5 mM EDTA, 0.1 mM PMSF) and lysed by sonication on ice for 2 minutes. Cell lysates were centrifuged at 15,000 rpm for 20 minutes. The clarified supernatant was applied to 150 μl of pre-equilibrated Amylose resin (NEB) in amylose buffer. After 1 hour incubation at 4 °C with slow rotation, the resin was transferred to disposable chromatography column, washed extensively with amylose buffer, and proteins were eluted with buffer containing 20 mM maltose. Eluted and input samples were resuspended in Laemmli buffer and analyzed by SDS–PAGE and Western blotting using an anti-His antibody (Cell Signaling Technology #2365). For pulldowns with *A. thaliana* and *S. pombe*, the pellet was resuspended with nickel buffer (25 mM HEPES-NaOH, pH 7.5, 500 mM NaCl, 10% glycerol, 0.1 mM DTT, 40 mM imidazole, 0.1 mM PMSF), proteins were pulled down using NiNTA resin (Thermo Scientific), eluted with buffer containing 500 mM imidazole, and analyzed by SDS–PAGE and Western blotting using anti-MBP antibody (NEB E8032).

#### Evolutionary analysis

Red1 and ASY3 homologs were identified through iterative sequence-similarity searches followed by reciprocal validation. Position-Specific Iterated BLAST (PSI-BLAST) searches were performed against the NCBI ClusteredNR protein database using fungal Red1 and plant ASY3 reference sequences as initial queries to identify candidate homologs in Fungi and Viridiplantae, respectively. Candidate proteins were evaluated by reciprocal best-hit searches against the corresponding reference proteomes; proteins that recovered the original Red1 or ASY3 query as the top reverse hit were retained as high-confidence homologs. To increase sensitivity and recover more divergent homologs, validated Red1- or ASY3-like sequences were then used as additional queries in successive rounds of PSI-BLAST searches. This iterative homolog-expansion procedure allowed newly validated homologs to serve as queries for detecting increasingly distant members of the Red1/ASY3 family. Candidates identified in later rounds were retained only when they showed reciprocal best-hit relationships with validated homologs and conserved Red1/ASY3-family features, as assessed by sequence alignment and AlphaFold3 structural prediction. Redundant entries, truncated fragments, and extreme length outliers were removed. When multiple entries represented the same species or closely related strain-level records, a single representative sequence was selected to reduce sampling bias while preserving phylogenetic breadth. The final curated datasets comprised 168 fungal Red1-family homologs and 248 plant ASY3 homologs (**Supplemental Datasets S1 and S2**).

For conservation analyses, curated homologs were partitioned into nested taxonomic datasets. Within each taxon-specific dataset, the corresponding reference sequence (*S. cerevisiae* Red1 or *A. thaliana* ASY3) was retained as the positional coordinate system, allowing conservation scores, alignment confidence, and domain or motif annotations to be compared residue by residue across taxonomic ranks. Multiple-sequence alignments were generated with MAFFT v7.511 using the L-INS-i algorithm (Katoh and Standley 2013), and column-wise alignment reliability was estimated with GUIDANCE2 (Sela et al. 2015).

Evolutionary conservation was quantified with ConSurf (Ashkenazy et al. 2016) using Bayesian Rate4Site inference (Pupko et al. 2002) for each taxon-specific multiple-sequence alignment. Substitution models were selected according to the protein and taxonomic rank. For Red1, JTT was used for Saccharomycetaceae, LG for Saccharomycotina, and WAG for Saccharomycetes, Ascomycota, and Fungi. For ASY3, JTT was used for Brassicaceae, Eudicotyledons, Magnoliopsida, and Streptophyta, and WAG for Viridiplantae. Reference positions represented by fewer than six ungapped sequences, or assigned ConSurf credibility intervals spanning four or more color grades, were classified as “no data” and were not interpreted. Conservation scores and GUIDANCE2 confidence values were mapped onto the reference proteins and displayed as residue-level heat maps together with curated domain and motif annotations.

### Data and code availability

Sequencing data have been deposited in the GEO database under accession numbers GSE337033 (ChIP-seq) and GSE337034 (S1-seq). Custom scripts for sequencing-data processing and the CellProfiler pipeline are available at https://github.com/HMurakami-lab. Amino acid sequences used for AlphaFold3 modeling are listed in **Supplemental Table S8**.

## Acknowledgements

We thank Bin Hu for plasmids, reagents, and access to laboratory facilities; Adèle Marston and Franz Klein for yeast strains; and Akira Shinohara for the anti-Rec8 antibody. We also thank Payal Gumber, Hana Oe, and Omar Nasr Ahmed Khalifa for experimental contributions; members of the Chromosome & Cellular Dynamics Section for reagents, instruments, and discussions; Alexander Lorenz for discussions and comments on the manuscript; and the Microscopy and Histology Core Facility at the University of Aberdeen for imaging support.

Author contributions: AR performed most experiments, data curation, formal analysis, computational analysis, visualization, and methodology development, and contributed to writing the original draft. YT performed initial ChIP-seq experiments and developed and performed calibrated S1-seq experiments and analyzed the data. TR performed the biochemical experiments. JUA performed the evolutionary conservation analysis and wrote the corresponding sections. HFA performed calibrated S1-seq experiments, wrote the corresponding Methods section, and provided experimental training and guidance to AR and YT. SH performed the primary AlphaFold structure prediction and contributed to manuscript review and editing. CCB contributed to conceptualization, supervision, resources, and manuscript review and editing. HM conceived and supervised the project, acquired funding, administered the project, performed ChIP-seq and S1-seq data analyses, and wrote the original draft with input from all authors.

This work was supported by the Medical Research Council Career Development Award (MR/W027313/1 to HM), and the Horizon Europe program of the European Research Council (CoG 101124379 to CCB). CCB is a Research Associate from the Fonds National de la Recherche Scientifique. AR was supported by the Elphinstone Scholarship and a School of Medicine award from the University of Aberdeen. YT was supported by the JST SPRING (JPMJSP2156) and JSPS KAKENHI (26KJ1875).

---

**Supplemental Table S1.** Spore viability patterns.

| Strain | 4:0 | 3:1 | 2:2 | 1:3 | 0:4 | Total tetrads | Spore viability (%) |
| --- | --- | --- | --- | --- | --- | --- | --- |
| wild type | 89 | 4 | 3 | 0 | 0 | 96 | 97.4 |
| *red1-ΔCIM* | 8 | 0 | 6 | 3 | 103 | 120 | 9.8 |
| *red1-mCIM* | 30 | 3 | 30 | 10 | 47 | 120 | 41.5 |

**Supplemental Table S2.** Genetic distance and MI nondisjunction estimated by spore-autonomous fluorescence assay.

|  |  | wild type | *red1-ΔCIM* | *red1-mCIM* |
| --- | --- | --- | --- | --- |
| *CEN8-ARG4* | PD:TT:NPD | 1429:537:3 | 1082:133:1 | 1093:173:7 |
|  | cM ± SE | 14.09 ± 0.56 | 8.12 ± 0.91 | 8.44 ± 0.77 |
| *ARG4-THR1* | PD:TT:NPD | 1735:234:0 | 1140:86:0 | 1168:105:0 |
|  | cM ± SE | 5.94 ± 0.36 | 3.51 ± 0.36 | 4.12 ± 0.39 |
|  | MI-NDJ | 7/1976 | 722/1948 | 495/1768 |
|  |  | = 0.4% | = 37.1% | = 28.0% |
|  | (95% CI) | (0.1–0.7) | (34.9–39.3) | (25.9–30.1) |

The number of tetrads exhibiting fluorescent marker configurations in parental ditype (PD), tetratype (TT) and nonparental ditype (NPD) are shown. Genetic distances were calculated using Perkins equation [ $cM = 100 \times (6 \times NPD + TT) / (2 \times (PD + NPD + TT))$ ]. Standard errors (SE) were calculated using Stahl Lab Online Tools (https://elizabethhousworth.com/StahlLabOnlineTools/). Exact 95% confidence intervals were calculated using binomial tests in R.

**Supplemental Table S3.**
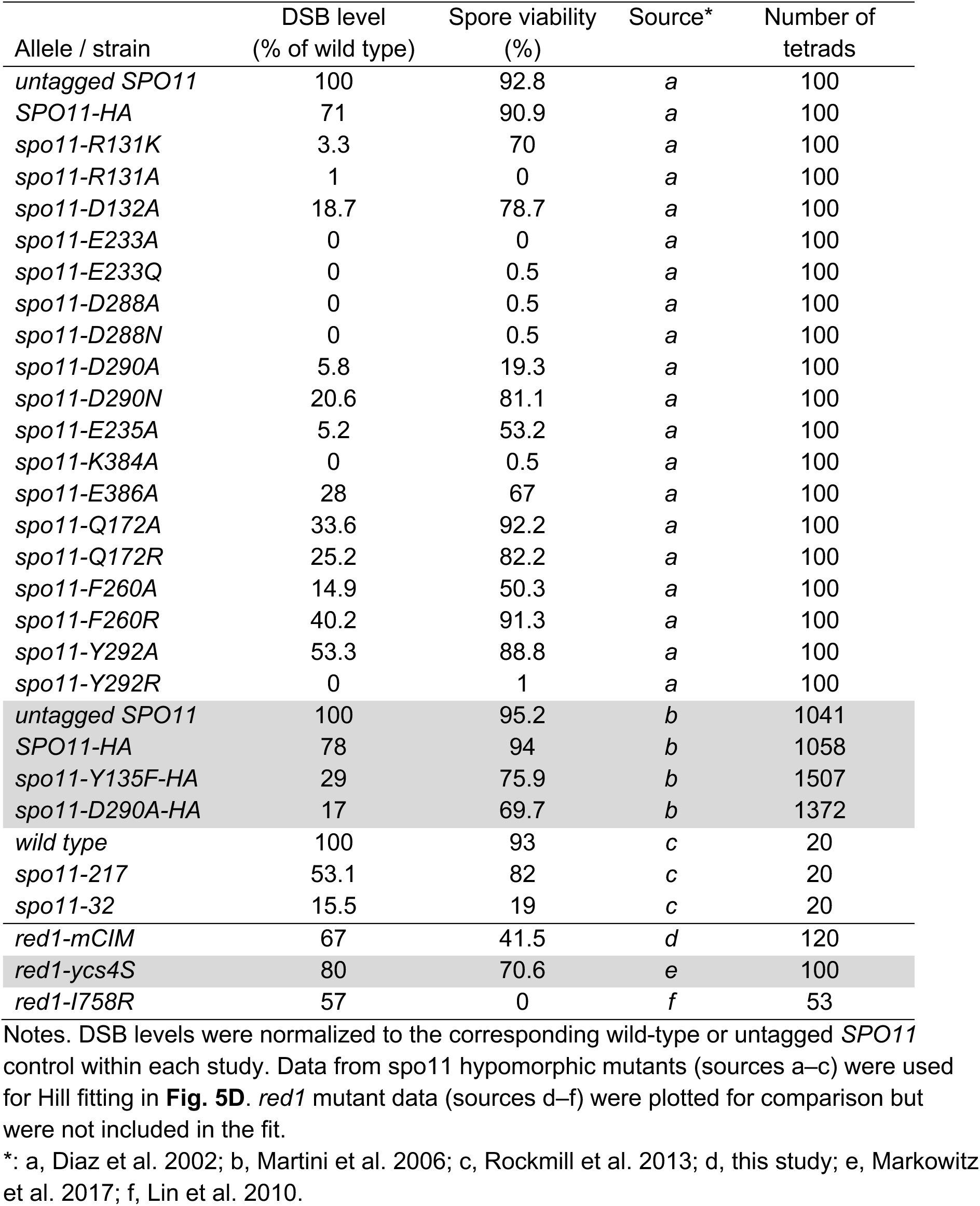
DSB levels and spore viability data used for DSB–spore viability modeling.

| Allele / strain | DSB level<br>(% of wild type) | Spore viability<br>(%) | Source* | Number of<br>tetrads |
| --- | --- | --- | --- | --- |
| *untagged SPO11* | 100 | 92.8 | *a* | 100 |
| *SPO11-HA* | 71 | 90.9 | *a* | 100 |
| *spo11-R131K* | 3.3 | 70 | *a* | 100 |
| *spo11-R131A* | 1 | 0 | *a* | 100 |
| *spo11-D132A* | 18.7 | 78.7 | *a* | 100 |
| *spo11-E233A* | 0 | 0 | *a* | 100 |
| *spo11-E233Q* | 0 | 0.5 | *a* | 100 |
| *spo11-D288A* | 0 | 0.5 | *a* | 100 |
| *spo11-D288N* | 0 | 0.5 | *a* | 100 |
| *spo11-D290A* | 5.8 | 19.3 | *a* | 100 |
| *spo11-D290N* | 20.6 | 81.1 | *a* | 100 |
| *spo11-E235A* | 5.2 | 53.2 | *a* | 100 |
| *spo11-K384A* | 0 | 0.5 | *a* | 100 |
| *spo11-E386A* | 28 | 67 | *a* | 100 |
| *spo11-Q172A* | 33.6 | 92.2 | *a* | 100 |
| *spo11-Q172R* | 25.2 | 82.2 | *a* | 100 |
| *spo11-F260A* | 14.9 | 50.3 | *a* | 100 |
| *spo11-F260R* | 40.2 | 91.3 | *a* | 100 |
| *spo11-Y292A* | 53.3 | 88.8 | *a* | 100 |
| *spo11-Y292R* | 0 | 1 | *a* | 100 |
| *untagged SPO11* | 100 | 95.2 | *b* | 1041 |
| *SPO11-HA* | 78 | 94 | *b* | 1058 |
| *spo11-Y135F-HA* | 29 | 75.9 | *b* | 1507 |
| *spo11-D290A-HA* | 17 | 69.7 | *b* | 1372 |
| *wild type* | 100 | 93 | *c* | 20 |
| *spo11-217* | 53.1 | 82 | *c* | 20 |
| *spo11-32* | 15.5 | 19 | *c* | 20 |
| *red1-mCIM* | 67 | 41.5 | *d* | 120 |
| *red1-ycs4S* | 80 | 70.6 | *e* | 100 |
| *red1-I758R* | 57 | 0 | *f* | 53 |

Notes. DSB levels were normalized to the corresponding wild-type or untagged *SPO11* control within each study. Data from *spo11* hypomorphic mutants (sources a–c) were used for Hill fitting in **Fig. 5D**. *red1* mutant data (sources d–f) were plotted for comparison but were not included in the fit.
\*: a, Diaz et al. 2002; b, Martini et al. 2006; c, Rockmill et al. 2013; d, this study; e, Markowitz et al. 2017; f, Lin et al. 2010.

**Supplemental Table S4.** Yeast strains used in this study.

| Strain number | Species | Genotype | Figures |
| --- | --- | --- | --- |
| **HMY892** | *S. cerevisiae* (SK1) | *MATa/α; ho::LYS2"/, lys2"/, ura3"/, leu2::hisG"/, his3::hisG"/, trp1::hisG"/, V5-RED1::loxP-KanMX-loxP"/, arg4-Nsp/arg4-Bgl* | 2A-G, 4A-D, 5A-B, S2A-B, S3A-D, S5A-B |
| **HMY894** | *S. cerevisiae* (SK1) | *MATa/α; ho::LYS2"/, lys2"/, ura3"/, leu2::hisG"/, his3::hisG"/, trp1::hisG"/, V5-red1(K553E,R554E,D555A,I558A,L559A,I562A,F563Y)::loxP-KanMX-loxP"/, arg4-Nsp/arg4-Bgl* | 2A-G, 4A-D, 5A-B, 5D, S2A-B, S3A-D, S5A-B |
| **HMY1135** | *S. cerevisiae* (SK1) | *MATa/α; ho::LYS2"/, lys2"/, ura3"/, leu2::hisG"/, his3::hisG"/, trp1::hisG"/, V5-red1(Δ553-563)::loxP-KanMX-loxP"/, arg4-Nsp/arg4-Bgl* | 2A-G, 5A-B, S2A-B, S2C S2D, S3A-D |
| **HMY390** | *S. mikatae* | *MATa/α; RAD50-V5::KanMX"/* | 2A-G, 4A-D, S3A-D |
| **HMY1656** | *S. cerevisiae* (SK1) | *MATa/α; ho::LYS2"/, lys2"/, ura3"/, leu2::hisG"/, his3::hisG"/, trp1::hisG"/, pREC8::SCC1-3HA::LEU2::rec8Δ::KanMX6"/, V5-RED1::loxP-KanMX-loxP"/* | 2A-G, S3A-D |
| **HMY1298** | *S. cerevisiae* (SK1) | *MATa/α; ho::LYS2"/, lys2"/, ura3"/, leu2::hisG"/, his3::hisG"/, trp1::hisG"/, sae2Δ::hphMX"/* | 3A-E, 4A-D, S4A-F, S5A-B |
| **HMY1300** | *S. cerevisiae* (SK1) | *MATa/α; ho::LYS2"/, lys2"/, ura3"/, leu2::hisG"/, his3::hisG"/, trp1::hisG"/, red1(K553E,R554E,D555A,I558A,L559A,I562A,F563Y)::loxP-KanMX-loxP"/, sae2Δ::hphMX"/* | 3A-E, 4A-D, S4A-F, S5A-B |
| **HMY1303** | *S. cerevisiae* (SK1) | *MATa/α; ho::LYS2"/, lys2"/, ura3"/, leu2::hisG"/, his3::hisG"/, trp1::hisG"/, red1Δ::CaURA3"/, sae2Δ::hphMX"/* | 3A-E, S4A-F |
| **HMY339** | *S. cerevisiae* (SK1) | *MATa/α; ho::LYS2"/, lys2"/, ura3"/, leu2::hisG"/, trp1::hisG"/, CEN8;ARG4;THR1::Cerulean-TRP1/CEN8::tdTomato-LEU2;ARG4::Yellow-URA3;THR1* | 5C |
| **HMY1154** | *S. cerevisiae* (SK1) | *MATa/α; ho::LYS2"/, lys2"/, ura3"/, leu2::hisG"/, his3::hisG"/, trp1::hisG"/, red1(Δ553-563)::loxP-KanMX-loxP"/, CEN8;ARG4;THR1::Cerulean-TRP1/CEN8::tdTomato-LEU2;ARG4::Yellow-URA3;THR1* | 5C |
| **HMY1159** | *S. cerevisiae* (SK1) | *MATa/α; ho::LYS2"/, lys2"/, ura3"/, leu2::hisG"/, his3::hisG"/, trp1::hisG"/, red1(K553E,R554E,D555A,I558A,L559A,I562A,F563Y)::loxP-KanMX-loxP"/, CEN8;ARG4;THR1::Cerulean-TRP1/CEN8::tdTomato-LEU2;ARG4::Yellow-URA3;THR1* | 5C |
| **HMY1232** | *S. cerevisiae* (SK1) | *MATa/α; ho::LYS2"/, lys2"/, ura3"/, leu2::hisG"/, his3::hisG"/, trp1::hisG"/, arg4-Nsp/arg4-Bgl* | 6A, S2C S2D |
| **HMY1233** | *S. cerevisiae*<br>(SK1) | *MATa/α; ho::LYS2"/, lys2"/, ura3"/,*<br>*leu2::hisG"/, his3::hisG"/, trp1::hisG"/,*<br>*red1(Δ553-563)::loxP-KanMX-loxP"/, arg4-*<br>*Nsp/arg4-Bgl* | 6A |
| **HMY1234** | *S. cerevisiae*<br>(SK1) | *MATa/α; ho::LYS2"/, lys2"/, ura3"/,*<br>*leu2::hisG"/, his3::hisG"/, trp1::hisG"/,*<br>*red1(K553E,R554E,D555A,I558A,L559A,I56*<br>*2A,F563Y)::loxP-KanMX-loxP"/, arg4-*<br>*Nsp/arg4-Bgl* | 6A |
| **HMY757** | *S. cerevisiae*<br>(SK1) | *MATa/α; ho::LYS2"/, lys2"/, ura3"/,*<br>*leu2::hisG"/, his3::hisG"/, trp1::hisG"/* | 6B, S2F |
| **HMY1448** | *S. cerevisiae*<br>(SK1) | *MATa/α; ho::LYS2"/, lys2"/, ura3"/,*<br>*leu2::hisG"/, his3::hisG"/, trp1::hisG"/,*<br>*red1(Δ553-563)::loxP-KanMX-loxP"/,*<br>*dmc1Δ::HphMX/"* | 6B |
| **HMY1449** | *S. cerevisiae*<br>(SK1) | *MATa/α; ho::LYS2"/, lys2"/, ura3"/,*<br>*leu2::hisG"/, his3::hisG"/, trp1::hisG"/,*<br>*red1(K553E,R554E,D555A,I558A,L559A,I56*<br>*2A,F563Y)::loxP-KanMX-loxP"/,*<br>*dmc1Δ::HphMX/"* | 6B |
| **HMY1452** | *S. cerevisiae*<br>(SK1) | *MATa/α; ho::LYS2"/, lys2"/, ura3"/,*<br>*leu2::hisG"/, his3::hisG"/, trp1::hisG"/,*<br>*dmc1Δ::HphMX/"* | 6B |
| **HMY888** | *S. cerevisiae*<br>(SK1) | *MATa/α; ho::LYS2"/, lys2"/, ura3"/,*<br>*leu2::hisG"/, his3::hisG"/, trp1::hisG"/, V5-*<br>*RED1::loxP-KanMX-loxP"/, arg4-Nsp/arg4-*<br>*Bgl* | S2C, S2D |
| **HMY890** | *S. cerevisiae*<br>(SK1) | *MATa/α; ho::LYS2"/, lys2"/, ura3"/,*<br>*leu2::hisG"/, his3::hisG"/, trp1::hisG"/, V5-*<br>*red1(K553E,R554E,D555A,I558A,L559A,I56*<br>*2A,F563Y)::loxP-KanMX-loxP"/, arg4-*<br>*Nsp/arg4-Bgl* | S2C, S2D |
| **HMY1372** | *S. mikatae* | *MATa/α; sae2Δ::KanMX/"* | 3A-E, 4A-D,<br>S4A-F, S5A-B |
| **HMY1670** | *S. cerevisiae*<br>(SK1) | *MATa/α; ho::LYS2"/, lys2"/, ura3"/,*<br>*leu2::hisG"/, his3::hisG"/, trp1::hisG"/,*<br>*pREC8::rec8(F581A)::KITRP1::tREC8(W303)*<br>*/"* | S2F |
| **HMY1672** | *S. cerevisiae*<br>(SK1) | *MATa/α; ho::LYS2"/, lys2"/, ura3"/,*<br>*leu2::hisG"/, his3::hisG"/, trp1::hisG"/,*<br>*pREC8::rec8(Y584A)::KITRP1::tREC8(W303)*<br>*/"* | S2F |
| **HMY1674** | *S. cerevisiae*<br>(SK1) | *MATa/α; ho::LYS2"/, lys2"/, ura3"/,*<br>*leu2::hisG"/, his3::hisG"/, trp1::hisG"/,*<br>*pREC8::rec8(P613K)::KITRP1::tREC8(W303)*<br>*/"* | S2F |
| **HMY1583** | *S. cerevisiae*<br>(SK1) | *MATa/α; ho::LYS2"/, lys2"/, ura3"/,*<br>*leu2::hisG"/, his3::hisG"/, trp1::hisG"/,*<br>*pREC8::rec8-red1(543-*<br>*564)::KITRP1::tREC8(W303)/"* | S2F |
| **HMY418** | *S. cerevisiae*<br>(SK1) | *MATa/α; ho::LYS2"/, lys2"/, ura3"/,*<br>*leu2::hisG"/, his3::hisG"/, trp1::hisG"/,*<br>*rec8Δ::KanMX6"/, arg4-Nsp/arg4-Bgl* | S2F |

**Supplemental Table S5.** Plasmids used in this study.

(A) Plasmids used to generate yeast mutants
| Plasmid number | Construct | *E. coli* marker | Yeast marker | Restriction enzymes used (tagged; untagged) |
| --- | --- | --- | --- | --- |
| pHM27 | *pRED1-V5-RED1-loxP-KanMX-loxP-tRED1* | Amp | KanMX | NdeI/KspAI; BseJI/KspAI |
| pHM28 | *pRED1-V5-red1(Δ553–563)-loxP-KanMX-loxP-tRED1* | Amp | KanMX | NdeI/KspAI; BseJI/KspAI |
| pHM29 | *pRED1-V5-red1(K553E,R554E,D555A,I558A,L559A,I562A,F563Y)-loxP-KanMX-loxP-tRED1* | Amp | KanMX | NdeI/KspAI; BseJI/KspAI |
| pHM75 | *pREC8-rec8(F581A)-KITRP1-tREC8(W303)* | Amp | KITRP1 | MluI/EcoRV |
| pHM76 | *pREC8-rec8(Y584A)-KITRP1-tREC8(W303)* | Amp | KITRP1 | MluI/EcoRV |
| pHM64 | *pREC8-rec8(P613K)-KITRP1-tREC8(W303)* | Amp | KITRP1 | MluI/EcoRV |
| pHM65 | *pREC8-rec8-red1(543–564)-KITRP1-tREC8(W303)* | Amp | KITRP1 | MluI/BspTI |

(B) *E. coli* expression plasmids
| Plasmid number | HisSUMO-tagged protein | MBP-tagged protein | Substitutions |
| --- | --- | --- | --- |
| pCCB1114 | HisSUMO-ScRec8-CTD | MBP control | - |
| pCCB1116 | HisSUMO-SpRec8-CTD | MBP control | - |
| pCCB1122 | HisSUMO-AtSYN1-CTD | MBP control | - |
| pCCB1123 | HisSUMO-AtSYN1-CTD | MBP-AtASY3-CIM | - |
| pCCB1124 | HisSUMO-ScRec8-CTD | MBP-ScRed1-CIM | - |
| pCCB1125 | HisSUMO-SpRec8-CTD | MBP-SpRec10-CIM | - |
| pTR23 | HisSUMO-ScRec8-CTD | MBP-ScRed1-CIM | Red1 F563A |
| pTR24 | HisSUMO-ScRec8-CTD | MBP-ScRed1-CIM | Rec8 P613K |
| pTR25 | HisSUMO-ScRec8-CTD | MBP-ScRed1-CIM | Rec8 I636A |
| pTR26 | HisSUMO-ScRec8-CTD | MBP-ScRed1-CIM | Rec8 F581A |
| pTR27 | HisSUMO-ScRec8-CTD | MBP-ScRed1-CIM | Rec8 Y584A |
| pTR29 | HisSUMO-ScRec8-CTD-ScRed1-CIM fusion | MBP-ScRed1-CIM | - |
| pTR42 | HisSUMO-SpRec8-CTD | MBP-SpRec10-CIM | Rec10 W603A |
| pTR43 | HisSUMO-AtSYN1-CTD | MBP-AtASY3-CIM | ASY3 L196A, L200A, L204A |

Sc: *Saccharomyces cerevisiae*; Sp: *Schizosaccharomyces pombe*; At: *Arabidopsis thaliana*; CTD: C-terminal domain; CIM: cohesin-interacting motif.

**Supplemental Table S6.**
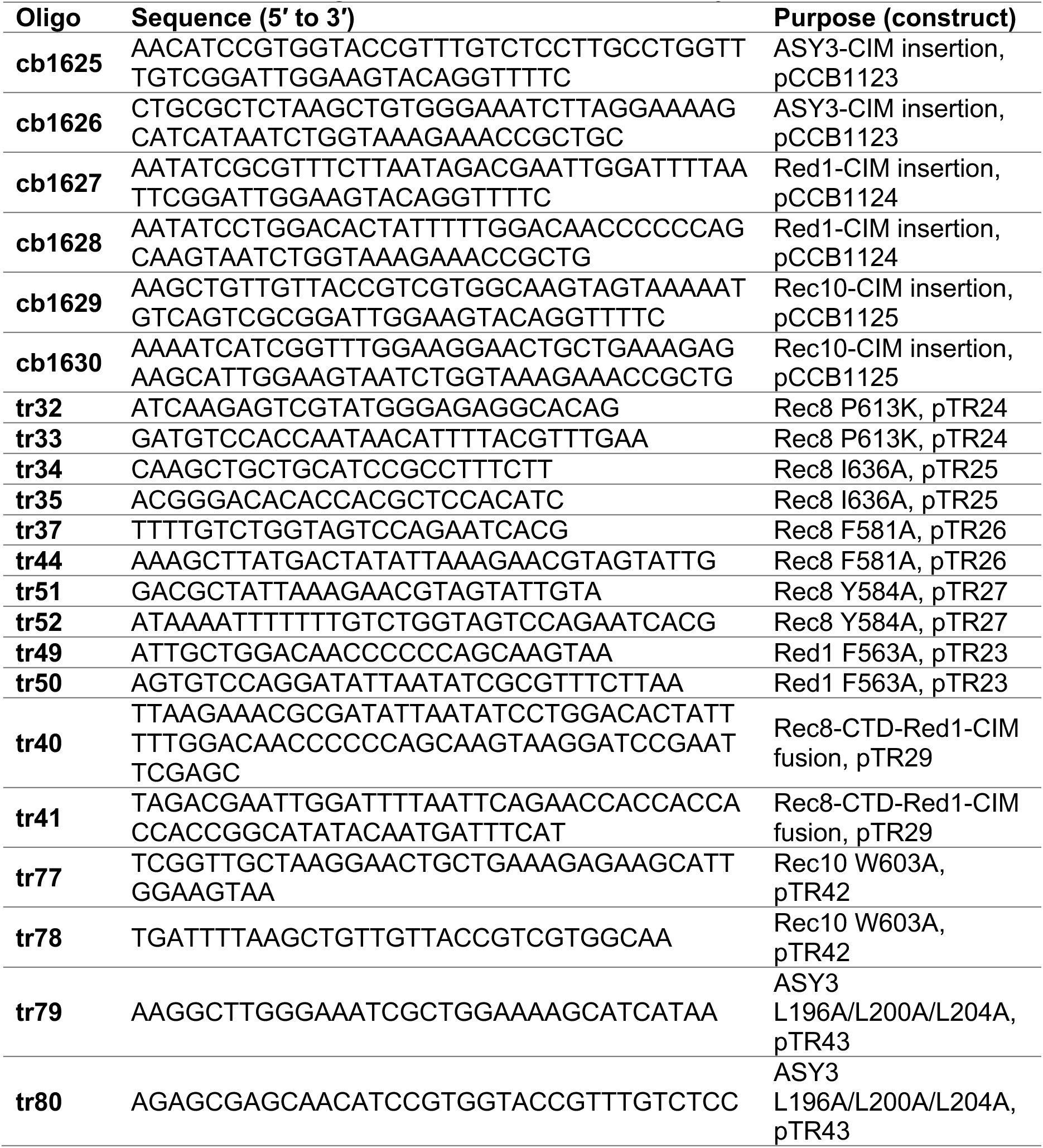
Oligonucleotides used in this study.

**Supplemental Table S7.**
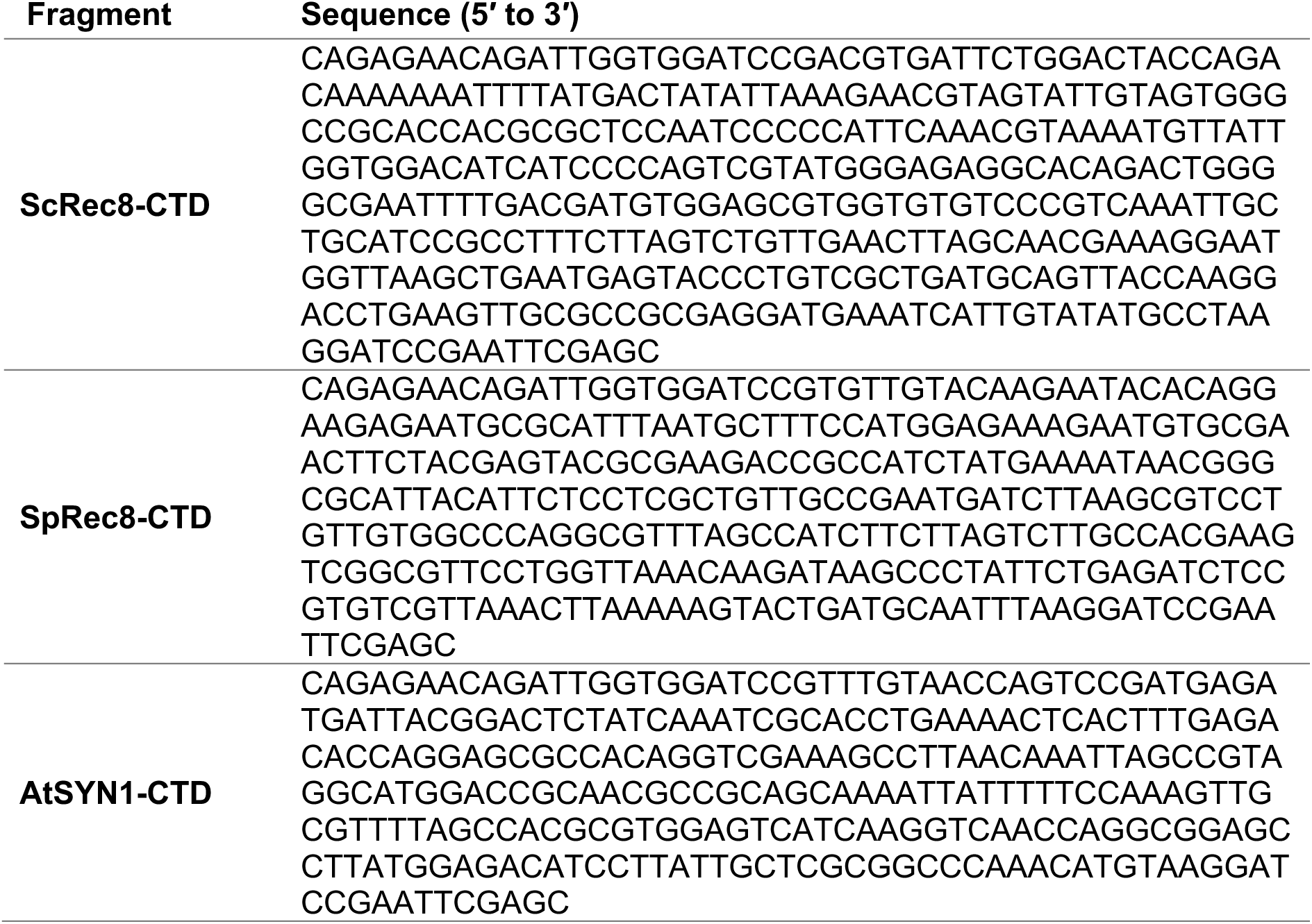
Synthetic DNA fragments (gBlocks) used in this study. Fragment Sequence (5′ to 3′)

**Supplemental Table S8.**
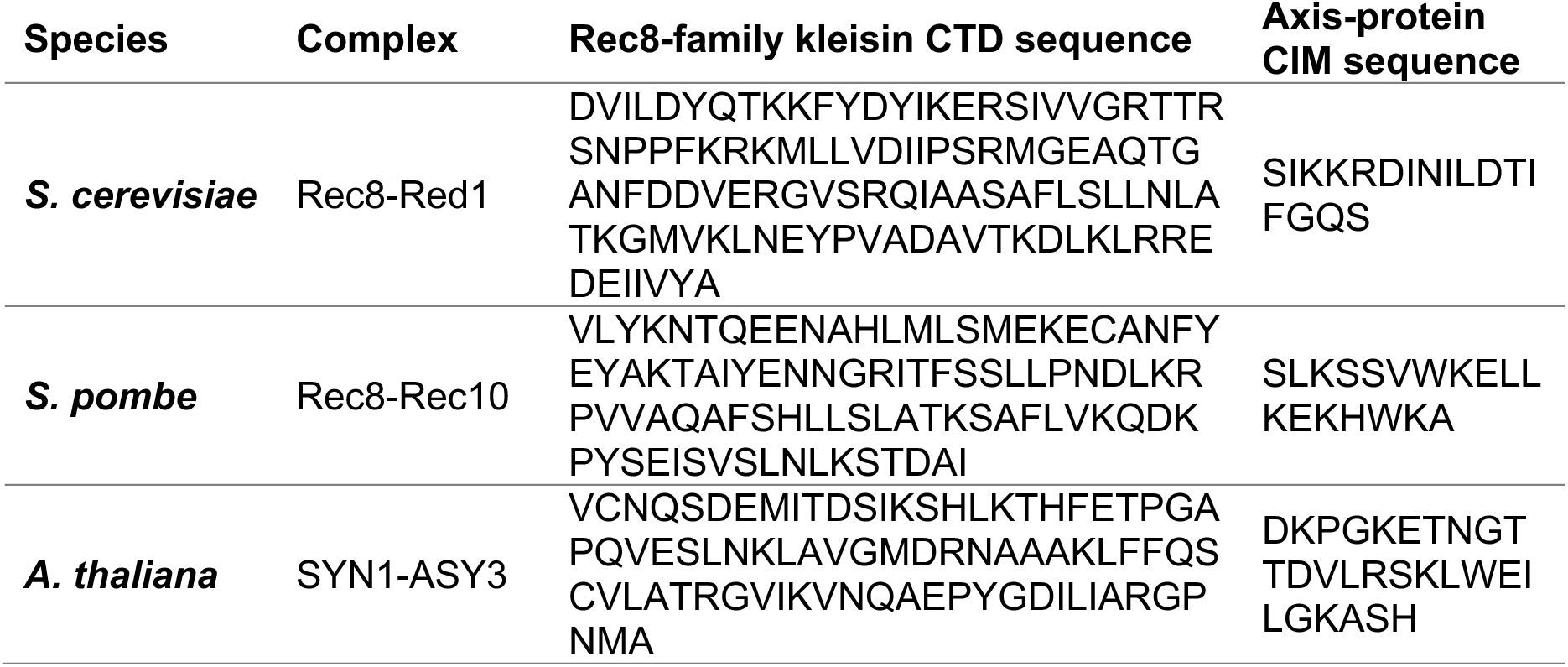
Protein sequences used as input for AlphaFold3 modeling.

**Supplemental Figure S1.**
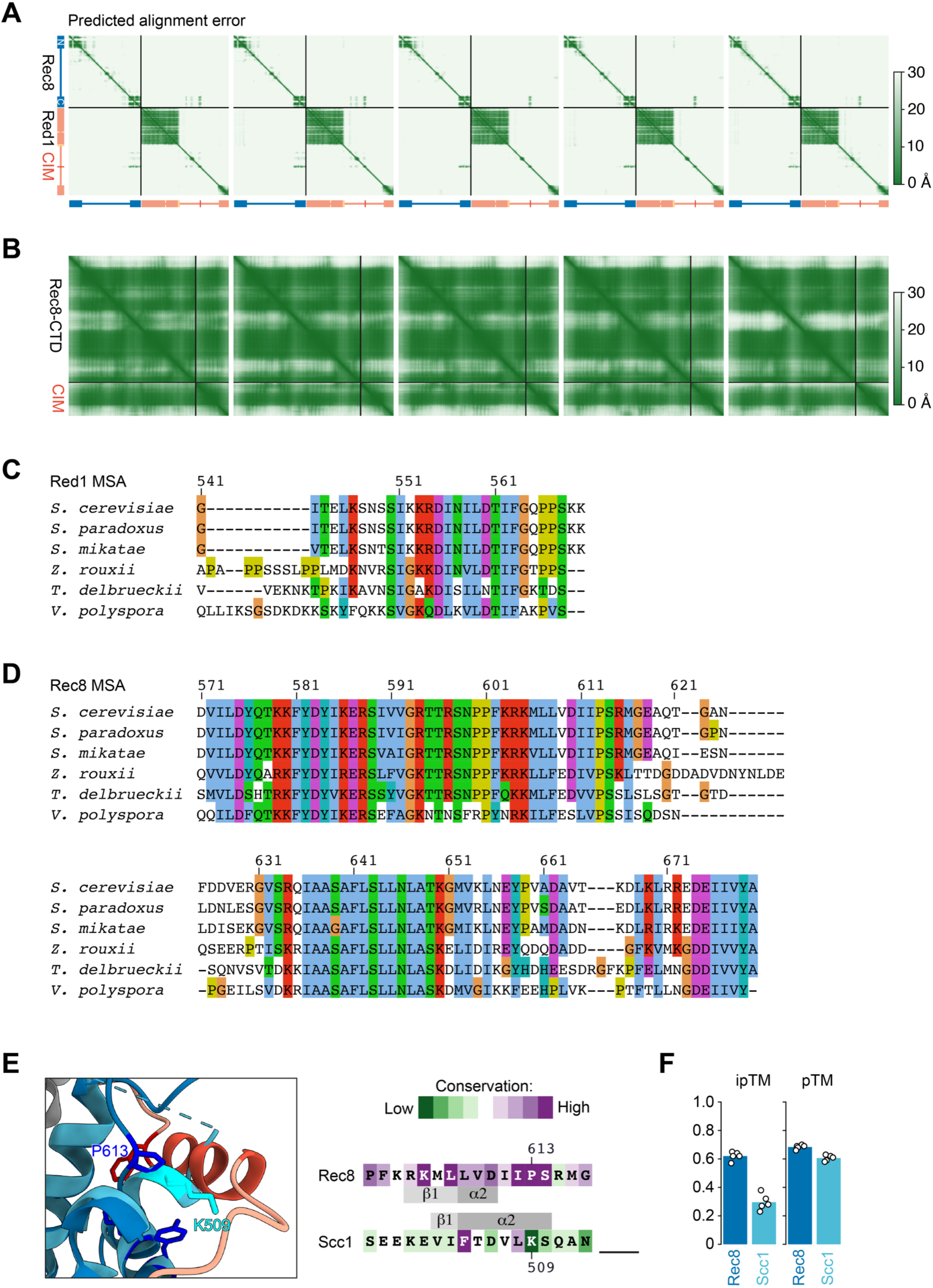
AlphaFold prediction and conservation analysis of the Red1–Rec8 interaction. **A.** AlphaFold3 prediction of the Red1–Rec8 interaction using full-length protein sequences. Predicted alignment error (PAE) plots support a well-defined interface between Red1-CIM and Rec8-CTD across independent predictions. **B.** AlphaFold3 modeling of the interaction using only the relevant regions of Red1 (543-573) and Rec8 (563-680). Global predicted template modeling score (pTM) and interface predicted template modeling score (ipTM) values for these models are shown in panel F. **C.** Multiple sequence alignment of the Red1 region containing the CIM across representative Saccharomycetaceae species. Numbers indicate positions in *S. cerevisiae*. **D.** Multiple sequence alignment of the Rec8-CTD. **E.** Structural comparison of the Rec8 CTD and the mitotic kleisin Scc1 (equivalent to **Fig. 1F**). Scc1 residues (504-510) form a longer helix than the equivalent one in Rec8, which may be incompatible with accommodating the Red1-CIM. **F.** AlphaFold3 modeling of the Red1 peptide with Scc1 (469-566). Under the same modeling conditions, no comparable interaction is predicted. Bars represent the mean of independent models (open circles).

**Supplemental Figure S2.**
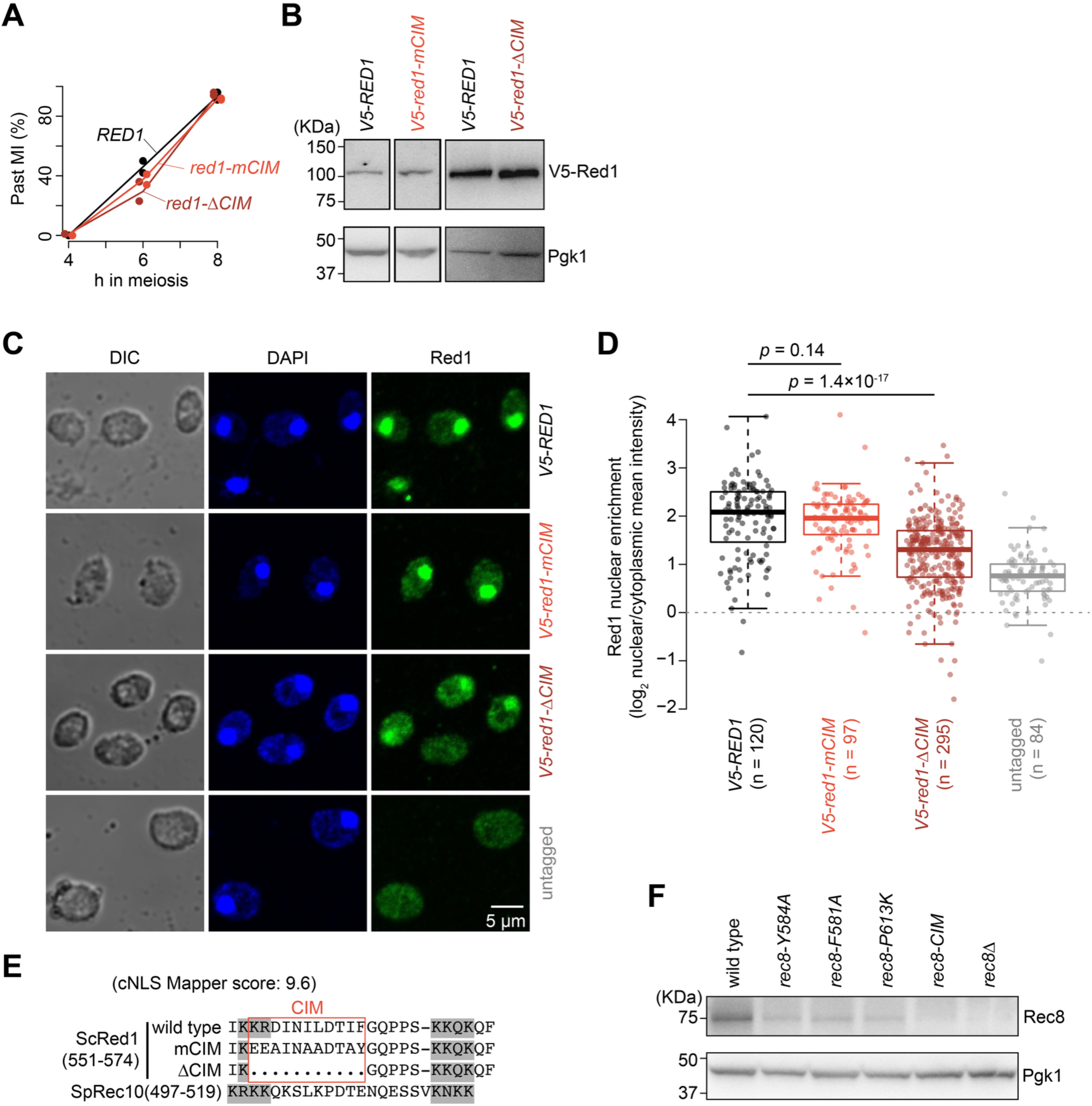
Characterization of *red1-CIM* mutant strains. **A.** Meiotic progression of wild type, *red1-ΔCIM*, and *red1-mCIM*. **B.** Immunoblot showing comparable V5-Red1 protein levels in wild type and CIM mutants. Cells were collected at 3 h after meiotic induction. **C.** Immunofluorescence of Red1 localization in meiotic nuclei (3 h in meiosis). **D.** Quantification of Red1 localization shown in panel C. The log2 ratio of nuclear/cytoplasmic mean Red1 signal (integrated signal / area) is plotted for individual nuclei. Solid and dashed lines indicate the mean and median, respectively. P values were calculated by two-sided Wilcoxon tests. In all box plots, thick horizontal bars denote medians, box edges mark the upper and lower quartiles, and whiskers indicate values within 1.5-fold of the interquartile range. **E.** Sequence comparison between the fission yeast Rec10 nuclear localization signal (NLS) and the corresponding region of budding yeast Red1. Residues shaded in grey indicate amino acids mutated in Rec10 to disrupt NLS function (Wintrebert et al. 2021). The Red1 sequence predicted as a bipartite NLS by cNLS Mapper (nls-mapper.iab.keio.ac.jp) is shown below, together with the mCIM and ΔCIM mutations. **F.** Immunoblot showing Rec8 protein levels in wild type and mutants detected by anti-Rec8 antibody (Bommi et al. 2019). Cells were collected at 3 h after meiotic induction.

**Supplemental Figure S3.**
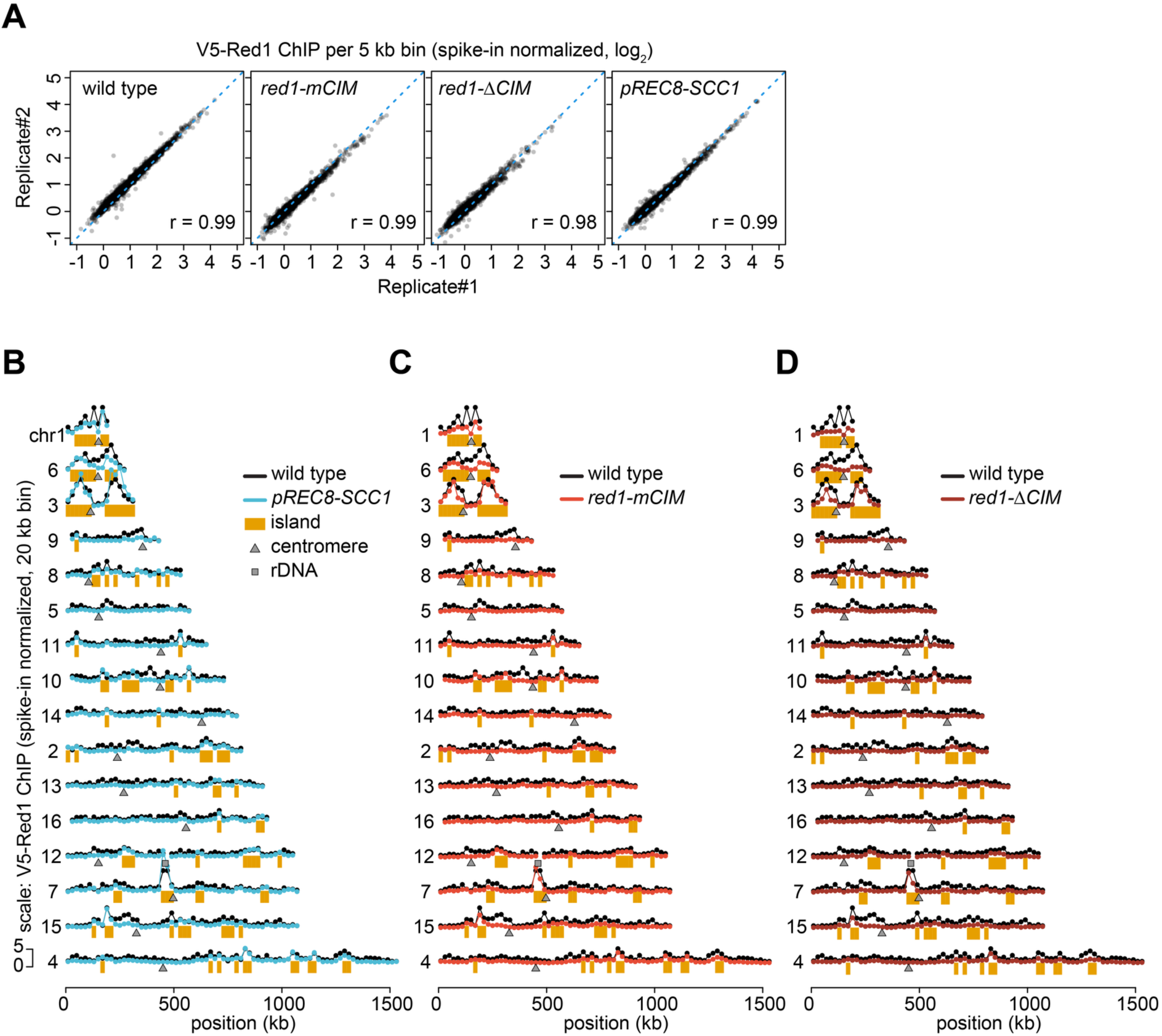
Reproducibility and genome-wide profiles of Red1 ChIP. **A.** Reproducibility between biological replicates of V5-Red1 ChIP-seq datasets. Spike-in normalized V5-Red1 ChIP-seq signals were calculated in 5-kb bins. Pearson correlation coefficients are indicated. **B.** Chromosome-wide V5-Red1 ChIP-seq profiles across all chromosomes in wild type and *pREC8-SCC1*. Signals were spike-in normalized and binned with a 20-kb window. Rec8-independent islands were defined using *pREC8-SCC1* as bins with Red1 ChIP signal greater than the mean + 0.3 SD; the remaining bins were classified as deserts. Centromeres and rDNA loci are indicated. **C.** Chromosome-wide Red1 ChIP profiles in wild type and *red1-mCIM*. **D.** Chromosome-wide Red1 ChIP profiles in wild type and *red1-ΔCIM*.

**Supplemental Figure S4.**
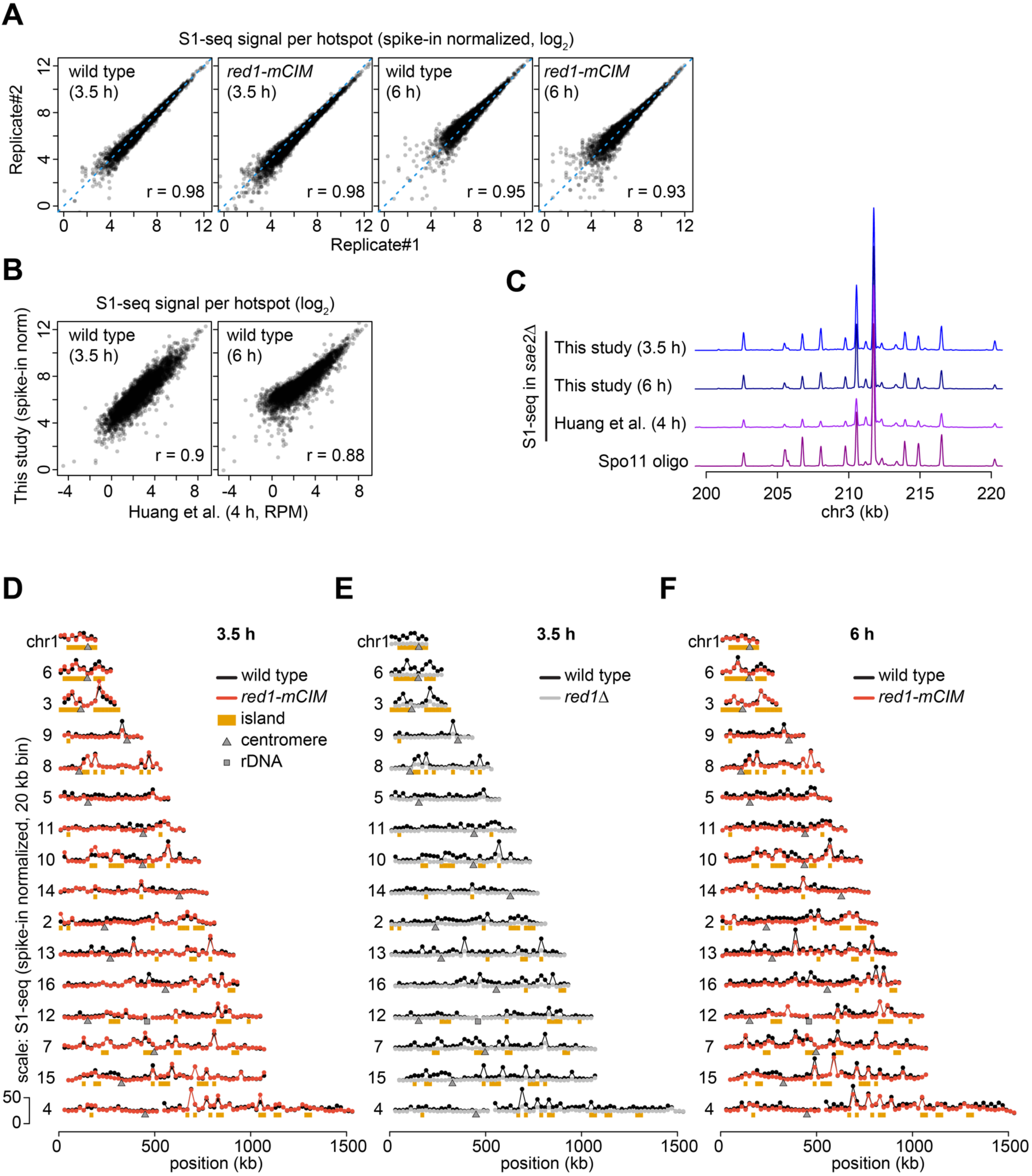
Reproducibility and genome-wide profiles of S1-seq. **A.** Reproducibility between biological replicates of S1-seq datasets. Spike-in normalized S1-seq signals were calculated for each hotspot in wild type and *red1-mCIM* at 3.5 h and 6 h after meiotic induction. Pearson correlation coefficients are indicated. **B.** Comparison of S1-seq hotspot signals generated in this study with a published S1-seq dataset. Hotspot-associated S1-seq signals in wild type at 3.5 h and 6 h were compared with the published 4 h dataset. Pearson correlation coefficients are indicated. **C.** Representative comparison of S1-seq profiles generated in this study with published S1-seq and Spo11-oligo profiles at chr3. Profiles from this study are shown for wild type at 3.5 h and 6 h after meiotic induction. **D.** Chromosome-wide S1-seq profiles across all chromosomes in wild type and *red1-mCIM* at 3.5 h after meiotic induction. Signals were spike-in normalized and calculated in 20-kb bins. Rec8-independent islands, centromeres, and rDNA loci are indicated. **E.** Chromosome-wide S1-seq profiles across all chromosomes in wild type and *red1Δ* at 3.5 h after meiotic induction. **F.** Chromosome-wide S1-seq profiles across all chromosomes in wild type and *red1-mCIM* at 6 h after meiotic induction.

**Supplemental Figure S5.**
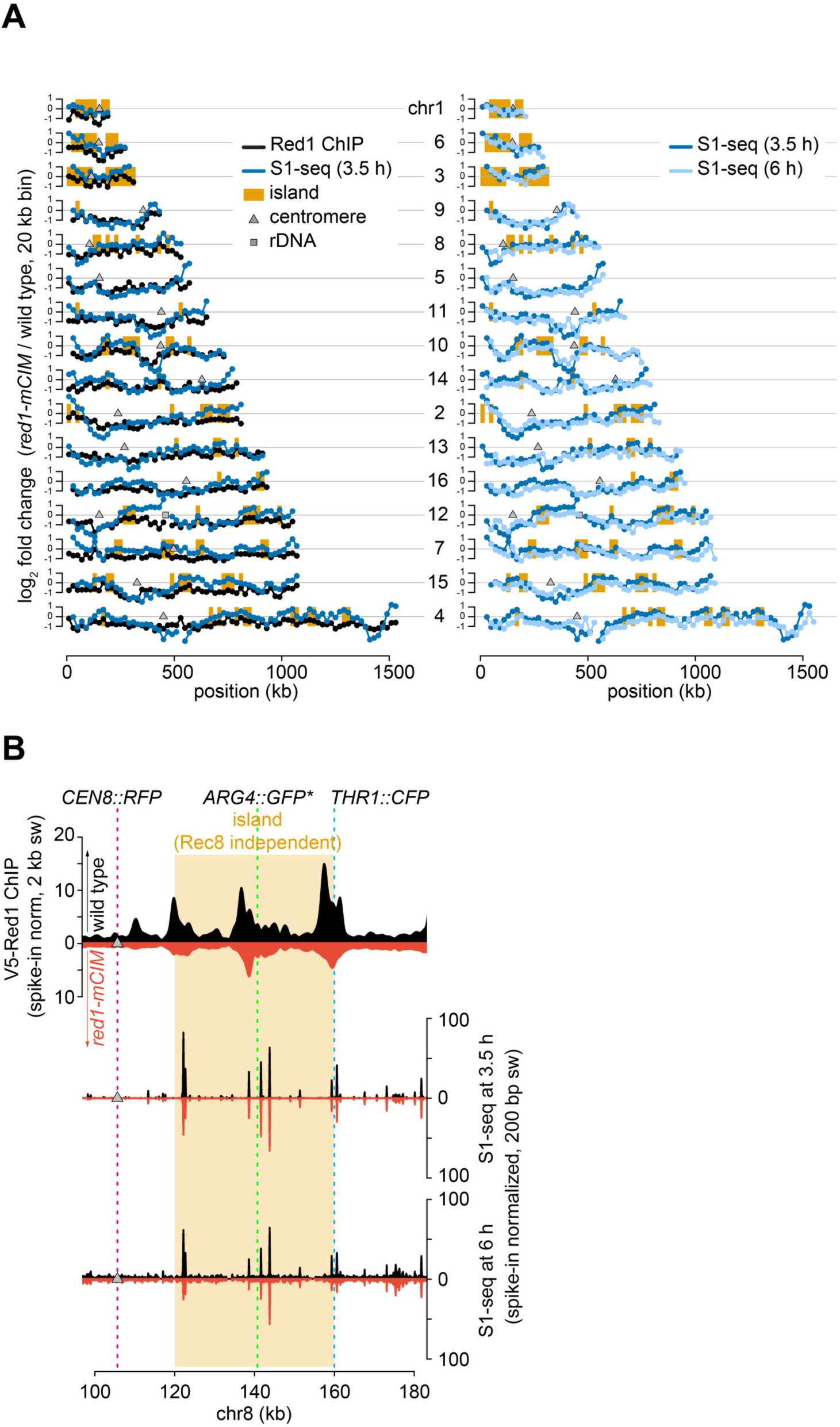
Genome-wide correspondence between Red1 ChIP and S1-seq changes in *red1-mCIM*. **A.** Chromosome-scale profiles of log2 fold changes in Red1 ChIP and S1-seq signal in *red1-mCIM* relative to wild type, calculated in 20-kb bins. Left: comparison of Red1 ChIP and S1-seq at 3.5 h. Right: comparison of S1-seq at 3.5 h and 6 h. Orange bars indicate island regions. Grey triangles and squares indicate centromeres and rDNA, respectively. **B.** Red1 ChIP and S1-seq profiles around the chr8 reporter interval used for genetic recombination and MI nondisjunction assays. The positions of *CEN8::RFP*, *ARG4::GFP*,* and *THR1::CFP* are shown as dashed lines. Wild-type signals are plotted above the baseline and *red1-mCIM* signals below the baseline. Red1 ChIP signals are shown as spike-in-normalized 2-kb Parzen sliding-window profiles. S1-seq signals at 3.5 h and 6 h are shown as spike-in-normalized 200-bp Parzen sliding-window profiles.

**Supplemental Figure S6.**
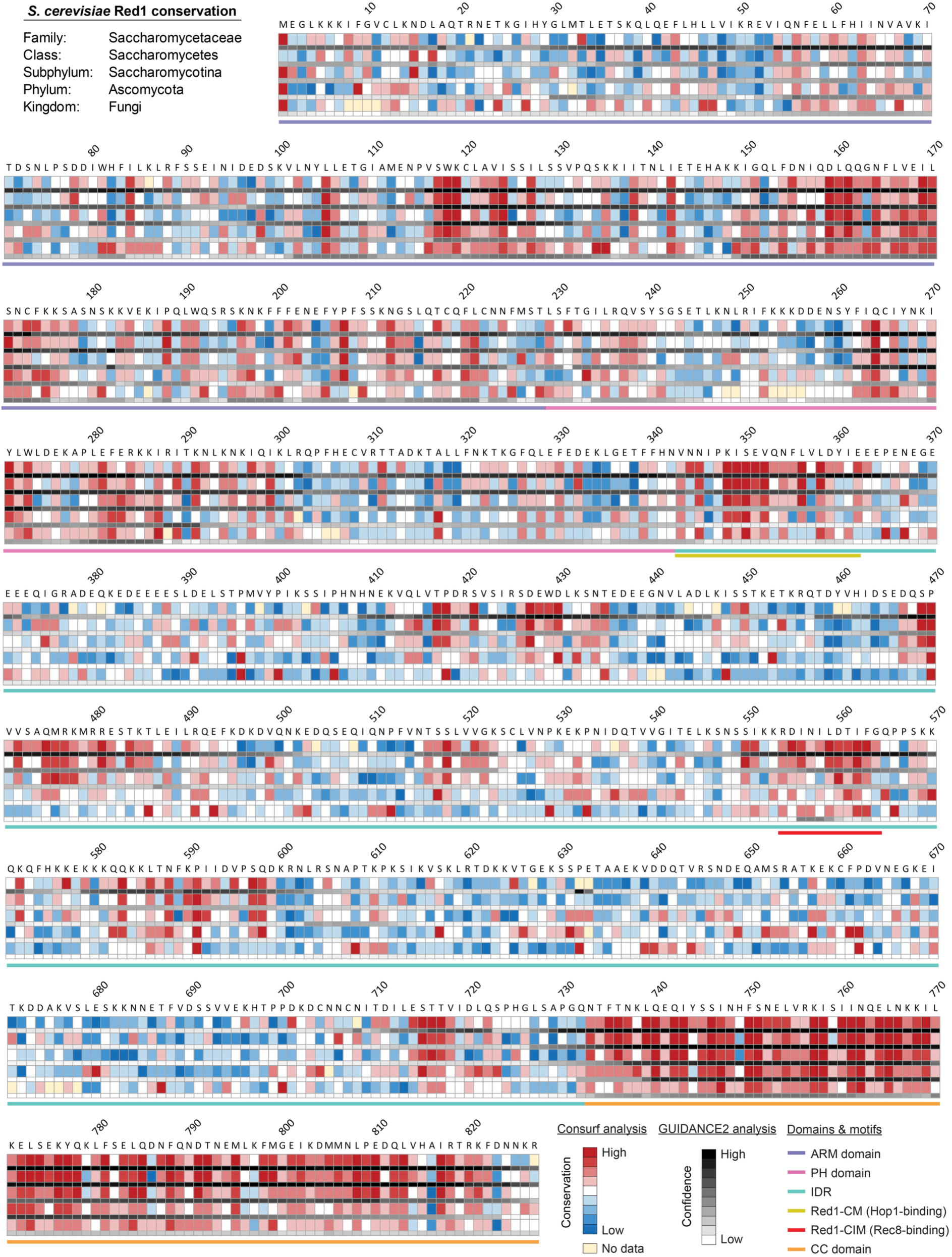
Conservation of *S. cerevisiae* Red1 across fungal taxonomic ranks. Residue–level conservation is mapped to the *S. cerevisiae* Red1 sequence across fungal taxonomic ranks. Heat maps report ConSurf scores and GUIDANCE2 confidence for **Supplemental Dataset S1** homologs. Annotated structural and functional features are indicated by colored bars below the sequence. ARM: armadillo repeat; PH: pleckstrin homology; IDR: intrinsically disordered region; CM: closure motif; CIM: cohesin-interacting motif; CC: coiled-coil.

**Supplemental Figure S7.**
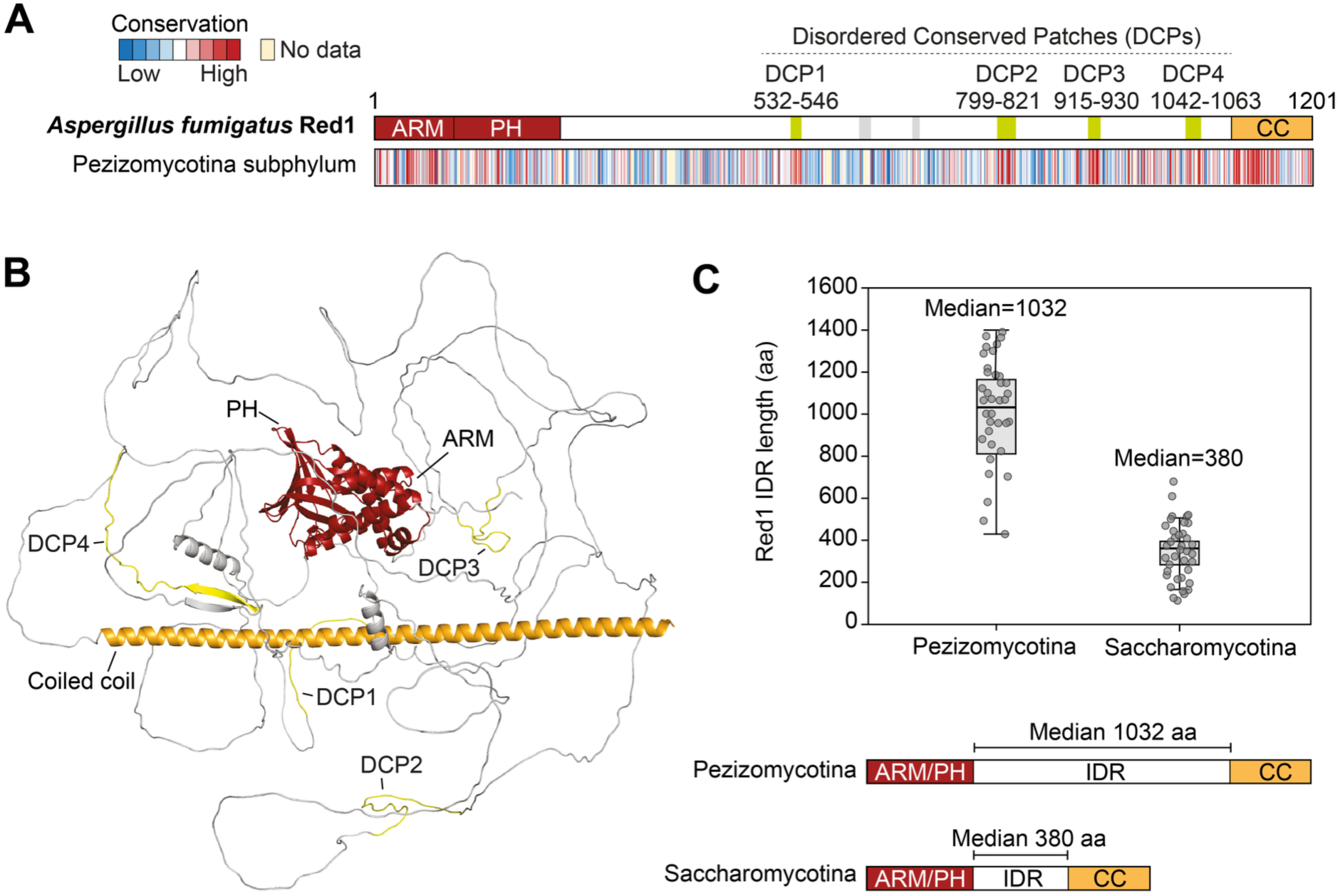
Expansion and divergence of the Red1 IDR in Pezizomycotina. **A.** Conservation profile of *Aspergillus fumigatus* Red1 across Pezizomycotina homologs from **Supplemental Dataset S1**. The domain organization of *A. fumigatus* Red1 is shown above the conservation track, with the ARM/PH domains, intrinsically disordered region (IDR), and C-terminal coiled-coil (CC) domain indicated. Disordered conserved patches (DCP1–DCP4) identified within the IDR are indicated in yellow. **B.** AlphaFold3 model of *A. fumigatus* Red1 (AF-B0XRH4-F1-v6), showing the ARM/PH domains, the C-terminal coiled-coil domain, and the four conserved patches within the expanded IDR. **C.** Distribution of predicted Red1 IDR lengths in Pezizomycotina and Saccharomycotina homologs. Each point represents one Red1 homolog; boxes indicate the interquartile range and horizontal lines indicate the median. Pezizomycotina Red1 homologs (n = 36) contain substantially longer IDRs than Saccharomycotina homologs (n = 38), with median lengths of 1032 aa and 380 aa, respectively. The schematic below illustrates the median-scaled Red1 domain organization in both subphyla.

**Supplemental Figure S8.**
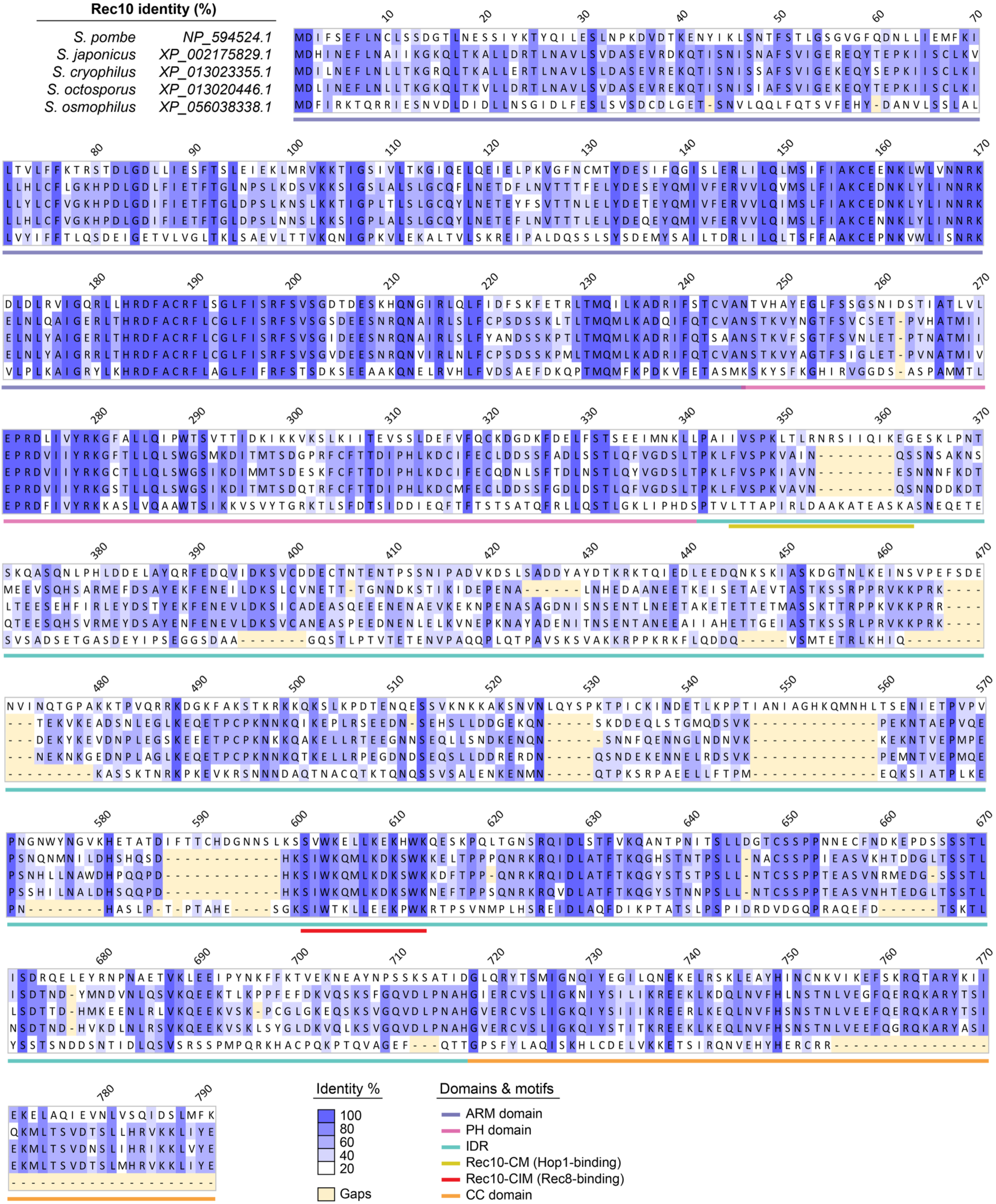
Conservation of *S. pombe* Rec10 across *Schizosaccharomyces* species. Multiple sequence alignment of Rec10 homologs from five fission-yeast species, trimmed relative to the ungapped *S. pombe* Rec10 reference sequence. Species names and NCBI accession numbers are shown on the left. Residue numbering corresponds to *S. pombe* Rec10. Column-wise sequence identity is represented by blue shading, with darker colors indicating higher conservation; gaps are shown in beige. Annotated structural and functional features are indicated by colored bars below the alignment. ARM, armadillo-repeat; PH, pleckstrin-homology; IDR, intrinsically disordered region; CM: closure motif; CIM: cohesin-interacting motif; CC: coiled-coil.

**Supplemental Figure S9.**
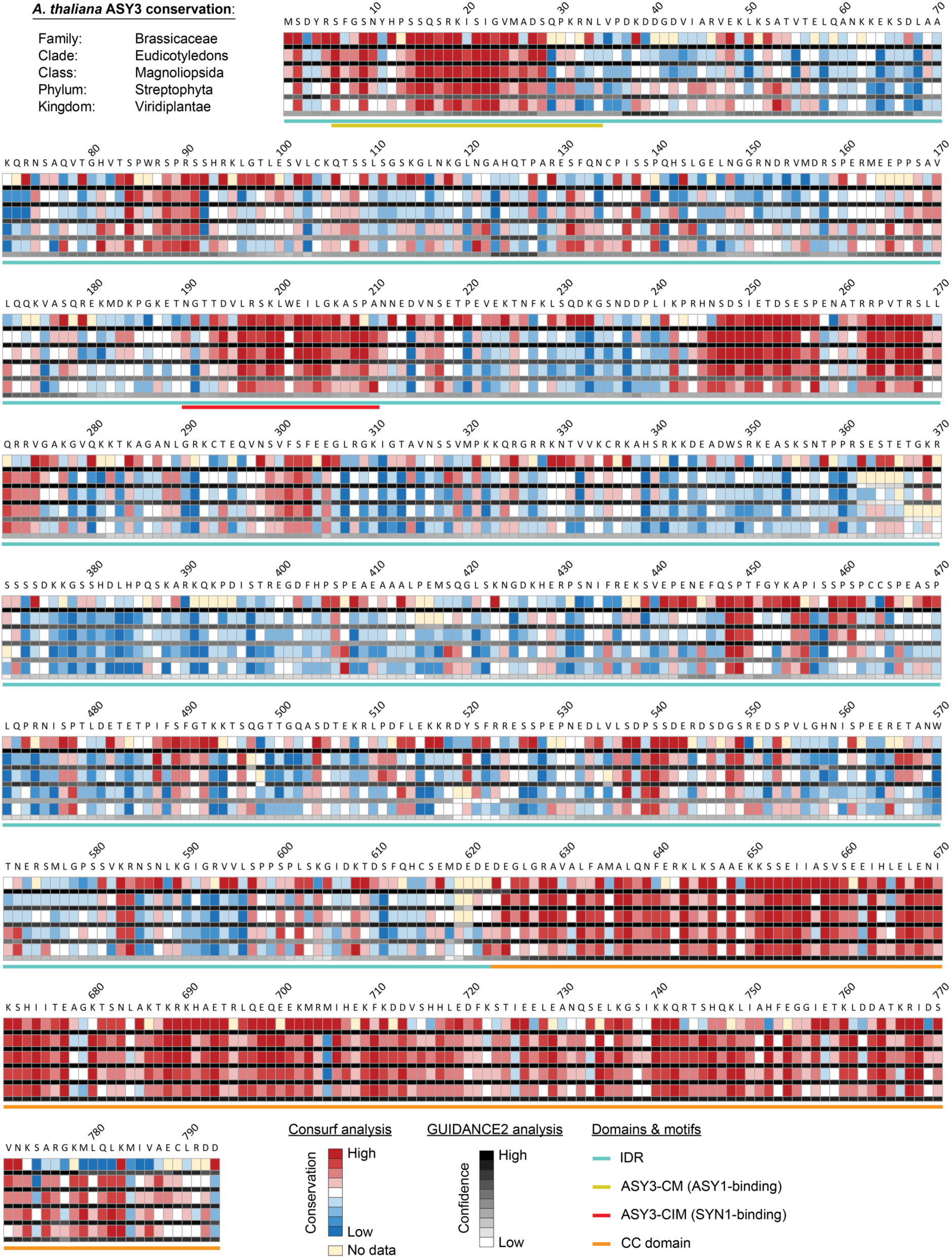
Conservation of *A. thaliana* ASY3 across plant taxonomic ranks. Residue-level conservation is mapped to the *A. thaliana* ASY3 sequence across nested plant taxonomic ranks. Heat maps report ConSurf scores and GUIDANCE2 confidence for **Supplemental Dataset S2** homologs. Annotated structural and functional features are indicated by colored bars below the sequence. IDR: intrinsically disordered region; CM: closure motif; CIM: cohesin-interacting motif; CC: coiled-coil.

